# Supraphysiological testosterone induces ferroptosis and activates NF-kappaB mediated immune pathways in prostate cancer through nucleophagy

**DOI:** 10.1101/2020.09.10.286252

**Authors:** Janet Mendonca, Rajendra Kumar, Olutosin Owoyemi, Kavya Boyapati, Naiju Thomas, Suthicha Kanacharoen, Max Coffey, Deven Topiwala, Carolina Gomes, Busra Ozbek, Tracy Jones, Marc Rosen, Liang Dong, Sadie Wiens, W. Nathaniel Brennen, John Isaacs, Angelo M. De Marzo, Mark Markowski, Emmanuel S. Antonarakis, David Qian, Kenneth Pienta, Drew Pardoll, Michael Carducci, Samuel Denmeade, Sushant Kachhap

## Abstract

The discovery that androgens play an important role in the progression of prostate cancer (PCa) has led to the development of androgen deprivation therapy as a first line of treatment against PCa. However, paradoxical growth inhibition has been observed, both experimentally and clinically, in a subset of PCa upon administration of supraphysiological levels of testosterone (SupraT). Here we report that SupraT activates cytoplasmic nucleic acid sensors and induces growth inhibition of SupraT-sensitive PCa cells. This is initiated by induction of two parallel autophagy-mediated processes, namely, ferritinophagy and nucleophagy. Consequently, autophagosomal DNA activates nucleic acid sensors that converge on NF-kappaB to drive immune signaling pathways. Chemokines and cytokines secreted by the tumor cells in response to SupraT results in increased migration of cytotoxic immune cells to tumor beds of animal xenografts and patient tumors. Collectively, our findings indicate that SupraT may inhibit a subset of PCa by activating nucleic acid sensors and downstream immune signaling.

## Main

In 1941, Charles Huggins discovered the benefits of androgen deprivation therapy (ADT), which has become the mainstay for advanced PCa treatment^1^. However, from the outset, it was recognized that all men eventually develop castration-resistant prostate cancer (CRPC)^2^. Evaluation of clinical specimens demonstrates that CRPC cells remain highly reliant on AR signaling^3,4^. These studies suggested that the adaptive reliance on AR signaling by CRPC cells becomes a therapeutic liability that can be exploited through the administration of SupraT, a concept we have termed bipolar androgen therapy (BAT)^5,6^. In this regard, we and others have demonstrated that the growth of some AR-positive human PCa cells can be inhibited by exposure to SupraT^6^. The mechanisms underlying this paradoxical effect of SupraT on prostate cancer cells are likely multifactorial as the androgen is the key mediator of prostate cancer cell metabolism, proliferation and death. Haffner *et al*. and others showed that androgens generate double-strand DNA breaks (DSBs) in PCa cells through the recruitment of AR and topoisomerase II beta to androgen response elements^7,8^. Hypothetically, in prostate tumors with DNA repair mutations, SupraT-induced DSBs would trigger DNA repair stress and lead to either growth inhibition or cell death^9^. Intuitively, these tumors would be acutely susceptible to BAT. In agreement with this postulate, we have recently discovered an association between germline and/or somatic DNA repair gene mutations and favorable response to BAT^10,11^. In this study, we report SupraT induces ferroptosis and nucleophagy-mediated immune activation resulting in growth inhibition of PCa.

Our previous work demonstrated that sensitivity to SupraT differed among PCa cell lines^6^. Cell cycle analysis revealed that SupraT causes a growth inhibitory accumulation of LNCaP cells in the G0/G1 phase, while LAPC4 cells show the opposite response and continue to proliferate (Extended Data Fig. 1a and b). Furthermore, clonogenic analysis revealed that SupraT significantly decreases the clonogenic potential of SupraT-sensitive LNCaP cells (Extended Data Fig. 1c and d). To understand whether sensitivity to SupraT reflected DNA repair mutations in these cancer lines, we searched the COSMIC cell line database for mutations in DNA repair genes in PCa lines (Extended Data Table 1). Strikingly, our analysis revealed that sensitivity to hormone exposure correlated directly with the number of DNA repair mutations harbored by the cell types (LNCaP > VCaP > LAPC4), mirroring our clinical data^6,11^. We hypothesized that due to the prevalence of DNA repair mutations, LNCaP cells treated with SupraT may not be able to repair DSBs induced by androgens and instead undergo DNA repair crisis leading to apoptosis. However, contrary to our hypothesis, LNCaP cells treated with synthetic androgen R1881, did not lead to a substantial increase in Annexin-V positivity or enhancement of PARP cleavage (Extended Data Fig. 2a and b). This suggested that the decrease in LNCaP cell number upon testosterone treatment may likely also involve a non-apoptotic cell death mechanism. We therefore probed whether necrotic markers were induced upon SupraT treatment (Extended Data Fig. 2c), and found them to be markedly reduced or unchanged, ruling out necrotic induction by SupraT. Recently, ferroptosis, a non-apoptotic cell death mechanism, has been linked to autophagy^12,13^. We first determined whether SupraT is able to induce autophagy in PCa lines. Treatment of PCa cell lines demonstrated that the synthetic androgen, R1881, notably induces autophagy in the SupraT sensitive LNCaP cells, while basal autophagy in the SupraT-insensitive LAPC4 cells remained unaffected upon treatment (Extended Data Fig. 3a). We did not find global upregulation of key autophagy proteins such as beclin or ATG12 (Extended Data Fig. 3b). However, we did find both the number of autophagosomes and autophagy flux (evaluated using an autophagy flux sensor) increased only in LNCaP cells upon treatment with androgens (Extended Data Fig. 3c). These results prompted us to determine whether SupraT is able to differentially induce ferroptosis in PCa cells that are sensitive to androgen.

Ferroptosis involves iron-dependent accumulation of toxic lipid peroxides that leads to cell death^14^. Degradation of the iron storage protein ferritin through a specialized form of autophagy, termed ferritinophagy, increases the labile pool of iron, leading to an increase in lipid peroxides^15^. We sought to determine whether ferritinophagy is induced by SupraT. As depicted in Fig. 1a, SupraT causes a dose-dependent decrease in ferritin levels in LNCaP cells as compared to LAPC4 cells, where ferritin levels remain unchanged. Confocal images of SupraT treated LNCaP cells revealed that ferritin colocalized with LC3B positive autophagosomes (Fig. 1b). Since NCOA4 interacts with ferritin and mediates its autophagic degradation^15-17^, we probed for NCOA4 and found that NCOA4 is induced by SupraT (Extended Data Fig. 4a), and interacts with ferritin in a SupraT dependent manner (Fig. 1c). Further, knockdown of NCOA4 in LNCaP cells inhibited R1881 induced ferritin degradation, and decreased cell death. (Fig. 1d & Extended Data Fig. 4b and c). To determine whether ferritin is indeed degraded through autophagy, we treated cells with hydroxychloroquine, an autophagy inhibitor, and MG-132, a proteasome inhibitor, and evaluated its effect on ferritin degradation upon SupraT treatment. As seen in Fig. 1e, hydroxychloroquine prevented the degradation of ferritin by SupraT, suggesting that ferritin is degraded through autophagy. We evaluated the functional consequence of decreased ferritin on lipid peroxide formation using the lipid peroxide sensor, C11-BODIPY^14,18^, which fluoresces green upon oxidation. As shown in Fig. 1f, R1881 treated LNCaP cells had increased lipid peroxides as compared to vehicle treated controls. We next investigated whether SupraT induces pro-ferroptotic gene expression^12,19^. Pro-ferroptotic genes such as *ALOX5, PTGS2*, and *NCOA4* were increased many folds over vehicle controls (Fig. 1g). We further confirmed that SupraT induced cell death in LNCaP cells is indeed through ferroptosis, as treatment with a combination of R1881 and a ferroptosis inhibitor, ferrostatin-1, abrogated cell death (Fig. 1h and i).

**Fig 1:**
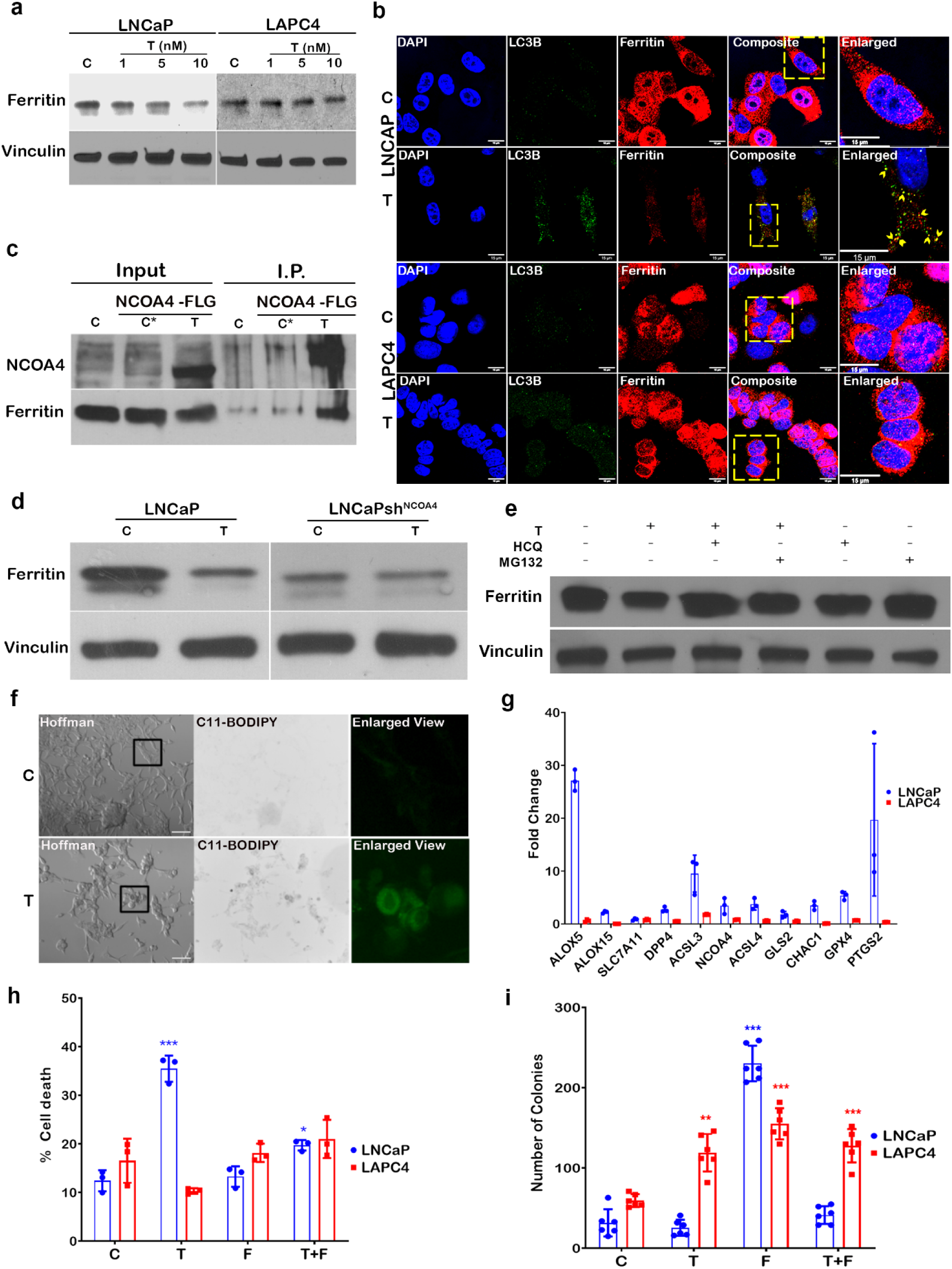
SupraT induces ferroptosis in PCa cells via induction of autophagy. **a**, Western blot analysis showing ferritin and loading control vinculin in LNCaP and LAPC4 cells treated with indicated concentrations (nM) of R1881 (T) and vehicle (C) for 72h. Representative of two independent experiments. Ferritin ∼ 21 kDa; loading control Vinculin 124 kDa. **b**, A representative single confocal section is shown from three independent experiments. LNCaP and LAPC4 cells stained for LC3B (green) and ferritin (red). Right most panels show enlarged inset images of a region of interest and arrow-head indicates colocalization of ferritin in autophagosomes (LC3B). **c**, Western blot for immunoprecipitated NCOA4-FLAG for interaction with ferritin in LNCaP cells transfected with human NCOA4α-pHAGE-C-FLAG-HA plasmid. Non-transfected (C) and NCOA4α-pHAGE-C-FLAG-HA transfected vehicle treated cells (C*) were used as controls. Representative of two independent experiments. NCOA4 70 kDa and Ferritin ∼ 21 kDa. **d**, Ferritin levels probed by western blot in vehicle (C) or R1881 (T) treated LNCaP or NCOA4 shRNA lentiviral particle transduced LNCaP cells. Representative of two independent experiments. Ferritin ∼ 21 kDa; loading control Vinculin 124 kDa. **e**, Western blot analysis for ferritin LNCaP cells treated with vehicle (C), T (10 nM R1881), HCQ (10 μM hydroxychloroquine), MG132 (10 μM) alone or combination of T with either of HCQ or MG132. Representative of two independent experiments. Ferritin ∼ 21 kDa; loading control Vinculin 124 kDa. **f**, A representative fluorescent microscopy images of control (C) and 10 nM R1881 treated (T) LNCaP cells from two independent experiments showing C11-BODIPY staining for oxidized membrane lipid (green fluorescence). Enlarged inset images in the rightmost panels show region of interest. **g**, Quantitative RT-PCR analysis for a panel of pro-ferroptotic genes in 10 nM R1881 treated LNCaP and LAPC4 cells post 72h. Vehicle control was used for normalization to calculate the fold change. Representative of three independent experiments. **h**, Cell death measured using trypan blue exclusion assay. Control (C), 10 nM R1881 (T), 10 µM Ferrostatin (F), or combination (T+F) shown as mean from three measurements with error bar showing standard deviation. **i**, Bar graph showing the mean of the number of colonies from six independent measurements counted using Fiji software for treatment conditions stated in (**h)** with error bars showing standard deviation. Asterisk indicates statistically significant differences (*p<0*.*05*). Scale bars: 15 µm (**b**) and 100 µm (**f**).

To investigate whether there is a direct link between response to SupraT, autophagy, and DNA repair, we treated SupraT sensitive LNCaP cells, which harbor several DNA repair gene mutations (Extended Data Table 1), with R1881 and stained cells for autophagosomes and DSBs. We postulated that cells harboring damaged DNA would have increased autophagy in response to DNA damage stress. Interestingly, LNCaP cells with increased autophagosomes displayed a smaller number of γ-H2AX puncta (Pearson’s correlation = −0.78) (Fig. 2a and b). This suggested that cells undergoing autophagy in response to SupraT might clear their damaged DNA more efficiently. Autophagy, being a dynamic event, renders visualization of fractions of autophagosomes carrying damaged DNA a challenge. Hydroxychloroquine prevents fusion of autophagosomes to lysosomes, leading to accumulation of autophagosomes^20^. Treatment of PCa cell lines with a combination of hydroxychloroquine and R1881 revealed marked localization of cytoplasmic DNA in LNCaP cells, as compared to the fewer cytoplasmic DNA puncta displayed in VCaP cells, which have intermediate DNA repair gene mutations (Extended Data Table 1). The SupraT-insensitive cells LAPC4, however, did not show any cytoplasmic DNA (Fig. 2c). To ensure that the observed cytoplasmic DNA was not mitochondrial DNA, we stained cells with antibodies against mitochondrial complex IV subunit I and LC3B. We did not find any notable co-localization of mitochondria with autophagosomes (Fig. 2d). However, we did find marked co-localization of cytoplasmic DAPI staining with LC3B in SupraT treated cells, indicating that the cytoplasmic DNA is present in autophagosomes (Fig. 2e). DAPI intensity peaked in the lumen of autophagosomes indicating that the DNA was indeed present within the autophagosomes (Fig. 2f). To ascertain whether the DNA within the autophagosomes harbored damaged DNA, we stained cells treated with a combination of hydroxychloroquine and SupraT with γ-H2AX and LC3B. As seen in Fig. 2g, LC3B and γ-H2AX co-localize with DAPI signals in the cytoplasm. This suggests that SupraT induced damaged DNA can be shuttled to the cytoplasm for autophagosome-mediated degradation. This implies that SupraT might induce two parallel autophagy-mediated phenomena: ferritinophagy and nucleophagy, both of which may be responsible for the growth inhibitory effects of SupraT.

**Fig 2:**
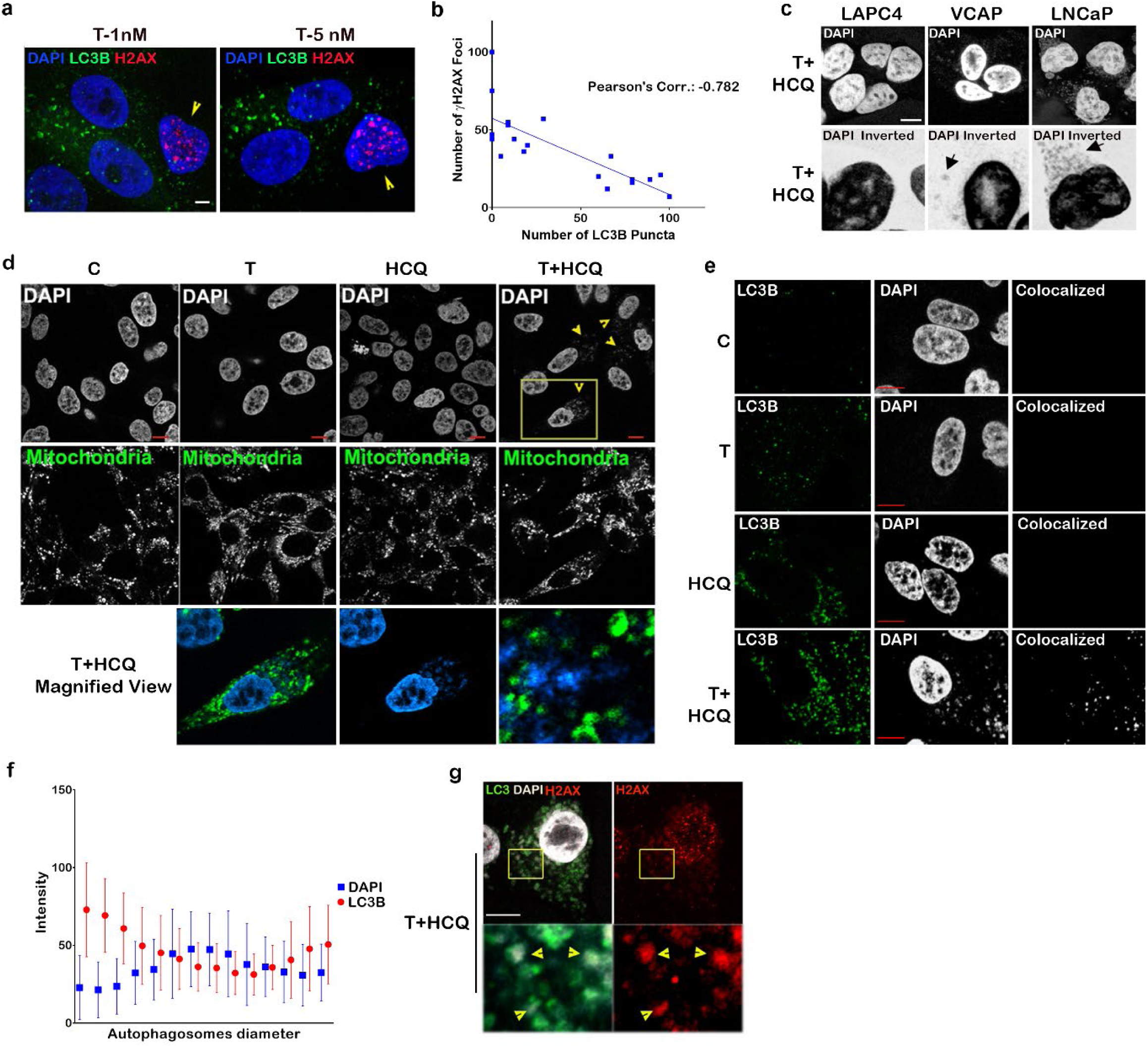
SupraT induces nucleophagic degradation of unrepaired damaged DNA. **a**, A representative single confocal section is shown from three independent experiments. LNCaP cells stained for γ-H2AX (red) and LC3B (green) after treatement with 1 and 5 nM R1881. Arrow-heads indicate cells harboring γ-H2AX foci with lower LC3B puncta. **b**, Scatter plot showing a correlation between the number of LC3B punctaand number of γ-H2AX foci. Counting of puncta was performed using the image analysis software Fiji. **c**, Confocal microscopy images for PCa cell lines treated with 10 nM R1881 (T) and hydroxychloroquine (HCQ). Lower panel is an inverted and magnified image of a single cell in the view-field for better visualization. Arrows indicate the localization of cytoplasmic DNA. **d**, Confocal images showing mitochondrial staining in LNCaP cells treated with vehicle (C), 10 nM R1881 (T), 10 µM HCQ or combination of T+HCQ for 72h. Lower panel shows a magnified view for enhanced visualization. **e**, Photomicrographs showing colocaliztion of LC3B (green) and DAPI (Gray) in LNCaP cells treated with control (C), 10 nM R1881 (T), HCQ (10 µM) or T+HCQ treated LNCaP cells (72h). Rightmost panel shows colocalized pixels. **f**, Graphical representation of fluorescence intensities of LC3B and DAPI on individual autophagosomes (n=16 measurements). **g**, Photomicrographs showing LC3B (green) and γ-H2AX (red) in LNCaP cells after 72h of treatment with T+HCQ. Lower panels show the enlarged insets with region of interest. Arrow-heads in lower-left image indicate presence of DNA (DAPI) in autophagosomes (LC3B) and lower-right image show γ-H2AX positivity in DNA present in those autophagosomes. Scale bars: 5µm (**a**), 10µm (**c**), 10µm (**d**), 10µm (**e**), and 10µm (**g**).

Cytosolic DNA is seen as a stimulant by the innate immune system as it is detected by DNA sensors in the cytoplasm that activate the adaptor protein STING and downstream innate immune signaling^21,22^. The STING promoter harbors an AR binding motif (Extended Data Fig. 5a), however, we did not find any induction of the STING transcript or STING promoter by treatment with SupraT (Extended Data Fig. 5b and c). Intriguingly, treatment with SupraT induced the STING protein as well as the RNA sensors RIG-I and MDA5 proteins in the SupraT sensitive LNCaP cells (Fig. 3a). Upon activation, monomeric STING dimerizes and translocates from the ER to autophagosome-like vesicles^23,24^. Separation of cellular homogenates on sucrose gradients revealed STING cofractionates with LC3B positive autophagosomes in SupraT treated cells (Fig. 3b). To confirm that STING is indeed activated upon SupraT treatment, we performed a STING dimerization assay, a gold standard for STING activation^22^. As shown in Fig. 3c, treatment with SupraT activates STING in LNCaP as compared to LAPC4 cells. Interestingly, SupraT also activated MAVS oligomerization indicating that the RNA sensing pathway was activated as well (Fig. 3d). Mislocalized and damaged DNA can also be detected by AIM2^25^. The AIM2 promoter harbors an AR binding motif (Extended Data Fig. 5d), hence we first measured AIM2 transcript levels. We did not find any induction of AIM2 promoter activity or AIM2 transcript levels by SupraT (Extended Data Fig. 5b and c). Neither did we find SupraT mediated activation of downstream inflammasomes, as evaluated by IL-1β cleavage, ruling out the involvement of AIM2 mediated inflammasome signaling by SupraT (Fig. 3f).

**Fig 3:**
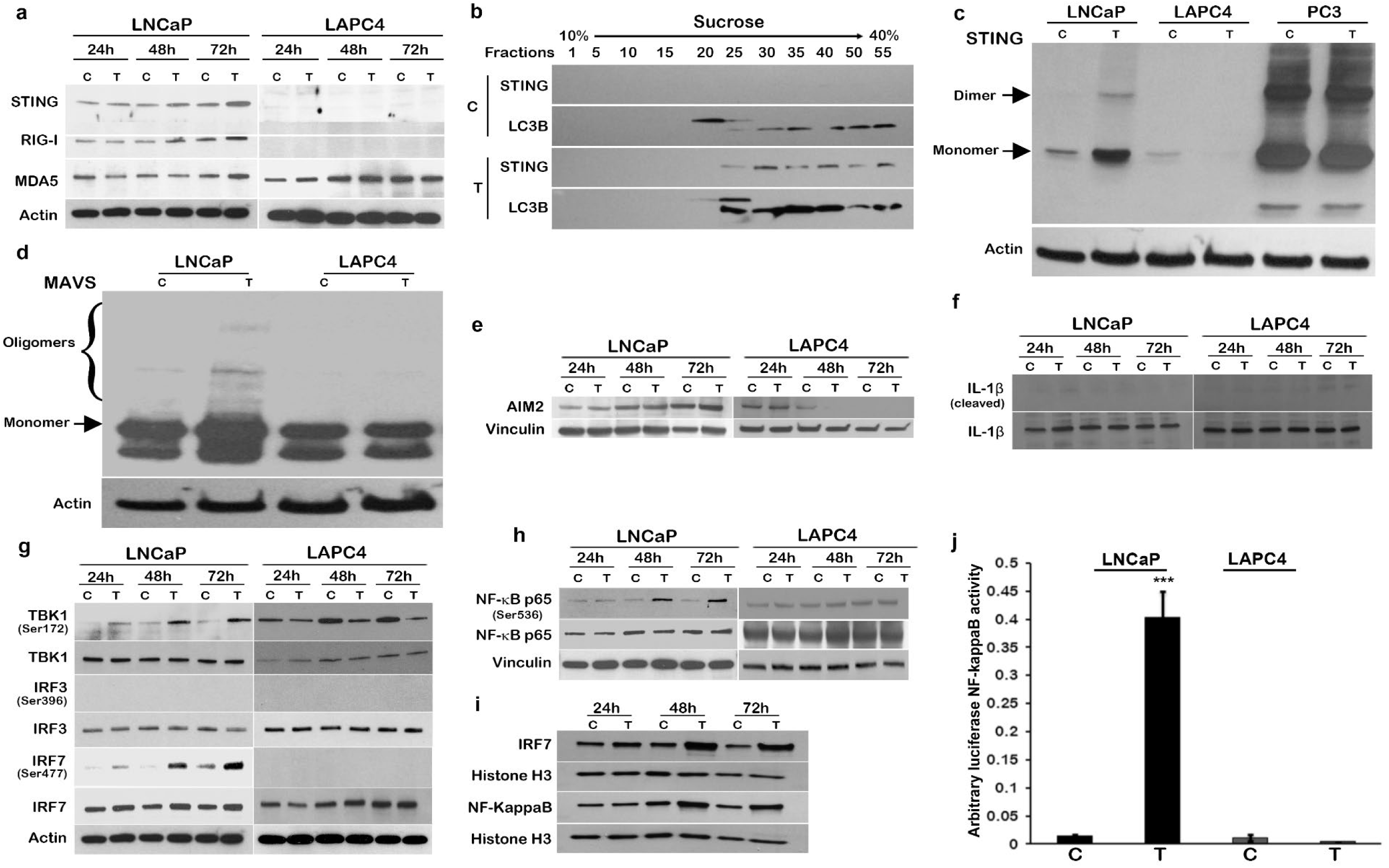
SupraT activates cytoplasmic nucleic acid sensors and subsequent NF-kappaB signaling. **a**, Western blot analysis showing DNA (STING) and RNA (RIG-I and MDA5) specific sensors along with actin as a loading control in LNCaP and LAPC4 cells treated with vehicle control (C) or 10 nM R1881 (T) in a time-dependent manner. Representative of two independent experiments. STING 33-35 kDa; RIG-I 102 kDa; MDA5 135 kDa; loading control Actin 43 kDa. **b**, Immunoblotting for LC3B and STING on fractions isolated using sucrose gradient centrifugation from LNCaP cell homogenates after 72h of 10 nM R1881 treatment with 10 nM R1881 (T) or vehicle control (C). Representative of two independent experiments. STING 33-35 kDa; LC3B 14, 16 kDa. **c-d**, Native western blot probing for STING **(c)** and MAVS **(d)** in PCa cell lines treated vehicle (C) or 10 nM R1881(T). Representative of two independent experiments; Loading control Actin 43 kDa. PC3 cells with constitutive expression of active STING were used as a positive control **(c). e-f**, Western blot analysis for AIM2 induction (AIM2 **(e)**) and signaling (IL-1β **(f)**) in PCa cell lines treated with vehicle (C) or 10 nM R1881 (T) in a time-dependent manner. Representative of two independent experiments. AIM2 40 kDa; IL-1β 17,31 kDa; Cleaved IL-1β 17 kDa; loading control Vinculin 124 kDa. **g-h** Immunoblotting for interferon regulatory genes (IRF7, IRF3, TBK1 **(g)**, and NF-kappaB **(h)**) in PCa cell lines treated with vehicle (C) or10 nM R1881 (T) at 24h, 48h, and 72h. Activated forms of all interferon regulatory genes were probed by respective phosphorylation specific antibodies. Representative of two independent experiments. Phospho-TBK1 84 kDa; TBK1 84 kDa; Phospho IRF3 45-55 kDa; IRF3 50-55 kDa, Phospho-IRF7 65 kDa; IRF7 65 kDa; loading control Actin 43 kDa. **i**, Nuclear localization of IRF-7 and NF-kappaB in LNCaP cells after treatment with vehicle (C) or 10 nM R1881 (T) at indicated time-points. Representative of two independent experiments. IRF7 65 kDa; NF-KappaB 65 kDa, loading control Histone H3 17 kDa. **j**, Bar graphs showing mean of luciferase NF-kappaB activity in LNCaP cells after treatment with vehicle (C) or 10 nM R1881 (T) using luciferase-based NF-kappaB reporter assay from three independent measurements and error bars show standard deviation. Asterisk indicates statistically significant differences (*p<0*.*05*).

Following its own activation, STING activates the Tank binding kinase 1 (TBK1), which in turn phosphorylates and activates interferon regulatory factors (IRFs), including IRF3, IRF7, and NF-kappaB, leading to the induction of immune response genes^21^. Although we did not find any activation of IRF3 by SupraT treatment (Extended Data Fig. 6 & Fig. 3g), our data indicates that TBK1, IRF7, and NF-kappaB canonical p65 subunit were activated upon SupraT treatment (Fig. 3g and h). Activated p65 and IRF7 translocated to the nucleus (Fig. 3i). We performed an NF-kappaB promoter reporter assay which confirmed the functional activation of NF-kappaB in SupraT sensitive cells (Fig. 3j). Intriguingly, we found SupraT stabilized NF-kappaB inducing kinase (NIK), a kinase central to the non-canonical NF-kappaB pathway (Fig. 4a). NIK phosphorylates the non-canonical p100 subunit of NF-kappaB and marks it for processing by a proteasome into the smaller p52 subunit, which subsequently dimerizes with RELB to activate transcription of target genes. Treatment with SupraT led to formation of p52 and induction of RELB in the SupraT sensitive cell line (Fig. 4a). Immunofluorescence confirmed nuclear translocation of p52 upon SupraT treatment (Fig. 4b). This data indicates that SupraT induced DNA damage activates nucleic acid sensors and downstream NF-kappaB signaling.

**Fig 4:**
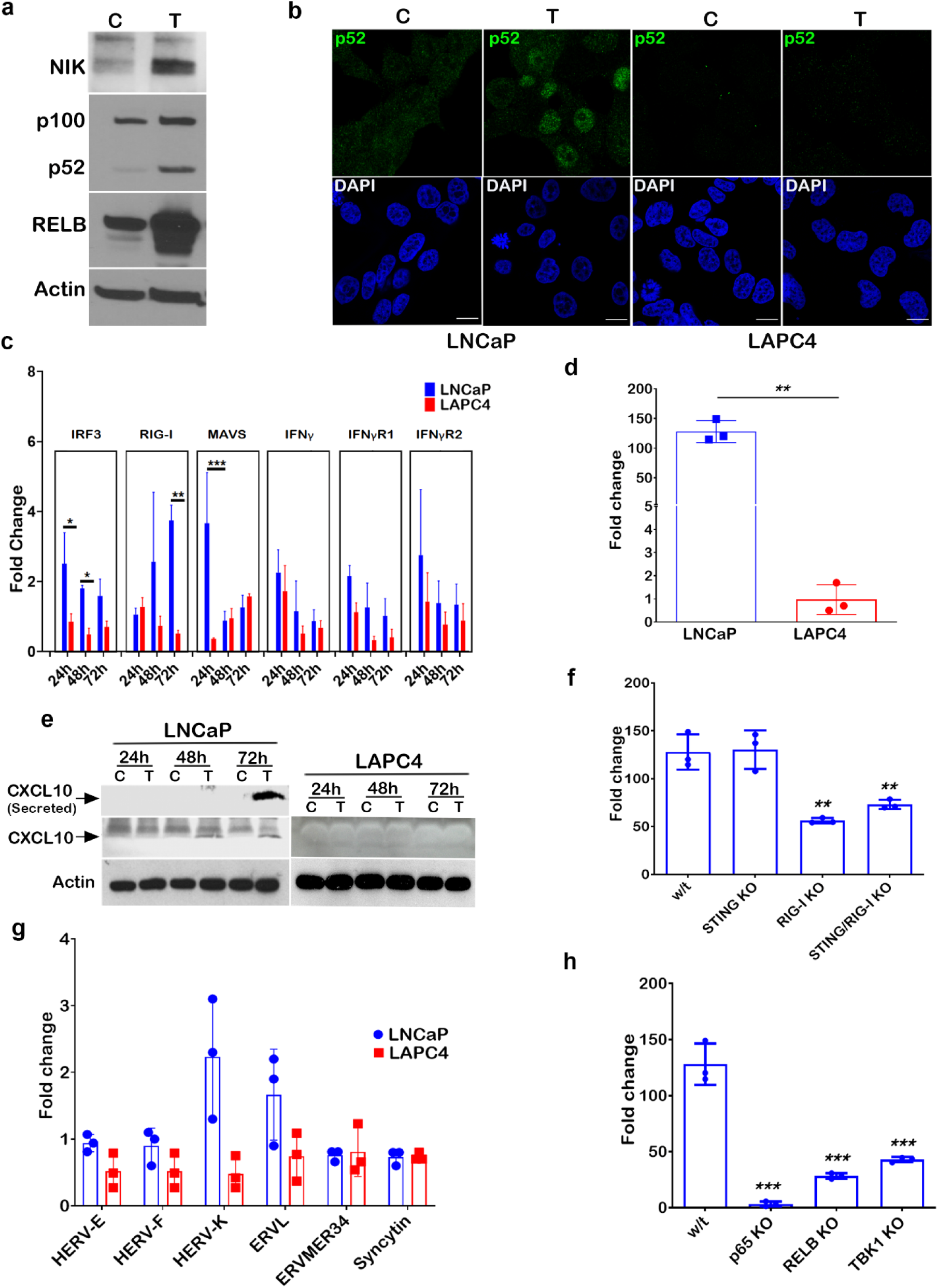
SupraT induced NF-kappaB signaling drives innate immune response. **a**, Immunoblotting for NIK, p100, p52, and RELB proteins in vehicle control (C) or 10 nM R1881 treated (T) LNCaP cells. Representative of two independent experiments. NIK 125 kDa; NF-kappaB 2 p100 and p52 120 and 52 kDa respectively; RELB 70 kDa; loading control Actin 43 kDa. **b**, Immunofluorescence images for the LNCaP and LAPC4 cells treated with vehicle (C) or 10 nM R1881 (T) and stained with p52 (green) and DAPI (blue) for visualization of nuclear p52 protein. Representative of two independent experiments. **c**, Quantitative RT-PCR for the interferon related genes in 10 nM R1881 treated LNCaP and LAPC4 cells for 24h, 48h, and 72h time points. Mean fold change data between LNCaP and LAPC4 cells were compared statistically, and an asterisk indicates a statistically significant difference (n=3). **d**, CXCL10 transcript analysis using quantitative RT-PCR from total RNA isolated from vehicle or 10 nM R1881 treated LNCaP and LAPC4 cells (n=3). **e**, Western blot for CXCL10 protein (lysates for cellular and culture media for secreted) for vehicle (C) and 10nM R1881 (T) treated LNCaP and LAPC4 cells made after 24h, 48h, or 72h post-treatment. Representative of two independent experiments. CXCL10 10 kDa; loading control Actin 43 kDa. **f**, RT-PCR based measurement for CXCL10 transcripts in LNCaP or knockouts of RIG-I, STING, or both after treatment with 10 nM R1881 for 72h. Mean fold change data (n=3) is plotted as histograms and asterisk indicates statistically significant difference (p<0.05). **g**, Histogram showing endogenous retroviral transcripts in 10 nMr R1881 treated LNCaP or LAPC4 cells. Bar indicates a mean fold change data (n=3), and the error bar indicates the standard deviation from three replicate values. **h**, Expression of CXCL10 transcripts in wild type, p65 KO, RELB KO, and TBK1 KO LNCaP cells treated with 10 nMr R1881. Statistical significance is calculated between the wild type and knockout variants of LNCaP cells. Mean fold change data (n=3) is plotted as histograms and asterisk indicates statistically significant difference (p<0.05). Scale bars: 15 µm (**b**).

In order to find out whether NF-kappaB responsive innate immune genes were activated by the nucleic acid sensors, we measured transcript levels of innate immune genes and found many genes were significantly upregulated in the SupraT-sensitive cell line (Fig. 4c). CXCL10, a chemokine, was increased^26^ several folds in treated LNCaP cells (Fig. 4d and e). Bioplex assays confirmed the induction and secretion of CXCL10 and other chemokines upon SupraT treatment (Supplementary Table 1). To determine whether STING or RIG-I sensors play a role in activation of downstream innate immune signaling, we created knockouts for STING, RIG-I, and STING/RIG-I double knockouts (Extended Data Fig. 7) and tested whether they abrogate CXCL10 induction. Knockout of STING did not abrogate CXCL10 induction but double knockouts and knockout of RIG-I alone decreased CXCL10 expression (Fig. 4f). This suggested that both the nucleic acid sensors are activated in SupraT-sensitive cells and that RIG-I may play an essential role in amplifying the signal. RIG-I can also be activated by the expression of endogenous retroviruses^27^. Our analysis revealed that some of the endogenous retroviruses harbor androgen response elements (Extended Data Fig. 8). However, none of the endogenous retroviral transcripts investigated were upregulated by SupraT (Fig. 4g), suggesting activation of RIG-I may be primarily through cytoplasmic DNA, as indicated by others^28^.

To tease out whether the canonical or the non-canonical NF-kappaB pathway is important for innate immune gene signaling by SupraT, we made knockouts of TBK1 and other key components of the NF-kappaB pathway (Extended Data Fig. 7). Knockouts of p65, TBK1, and RELB each abrogated downstream CXCL10 induction by SupraT (Fig. 4h). Induction or proteolytic processing of non-canonical subunits RELB and p100, respectively, by SupraT was totally abrogated in p65 and TBK1 knockouts (Fig. 5a), indicating that the canonical pathway is important for the activation of the non-canonical pathway, and TBK1 may play a critical role in linking the two. Furthermore, knockouts of p65, RELB, and TBK1 each abrogated NIK stabilization compared to parental cells, knockouts of STING and RIG-I decreased NIK stabilization, and STING/RIG-I double knockouts totally diminished the NIK stabilization, mirroring that of the p65 and TBK1 knockouts (Fig. 5b). Intriguingly, knockouts of STING did not revoke induction or processing of p100 but knockouts of RIG-I alone decreased p100 induction. Further, STING/RIG-I double knockouts had lower induction of RELB and p100 processing as compared to STING knockouts (Fig. 5c). While this data corroborates the involvement of both the nucleic acid sensors, STING and RIG-I may have opposite effects on the non-canonical NF-kappaB signaling induced by SupraT - STING being suppressive and RIG-I being supportive.

**Fig 5:**
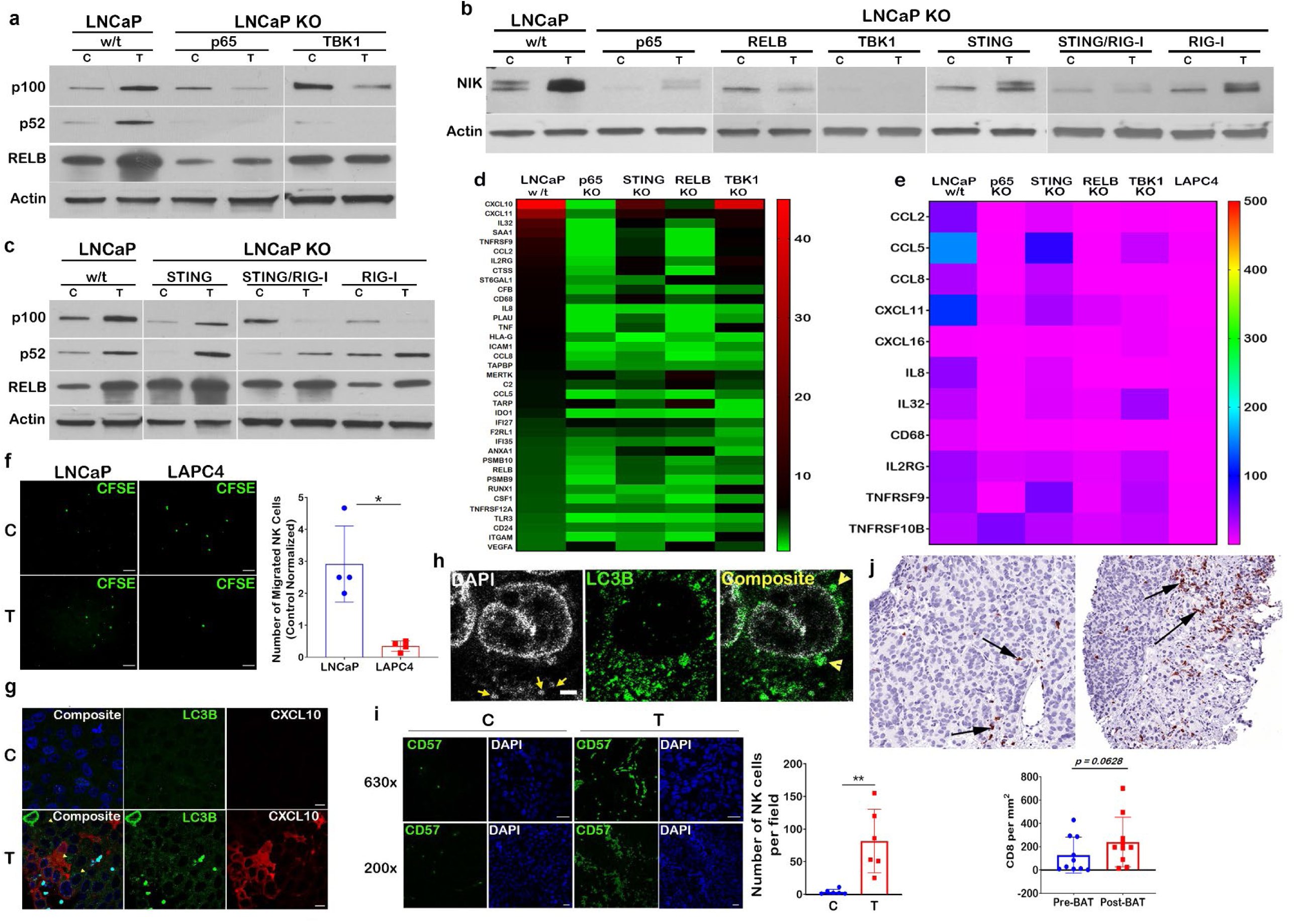
SupraT activates both canonical and non-canonical NF-kappaB signaling and induces tumor infiltration of immune cells. **a**, Immunoblotting for p100, p52, and RELB protein to show noncanonical NF-kappaB signaling in wild type and knockout LNCaP cells treated with vehicle (C) or 10 nM R1881 (T). Representative of two independent experiments. p100 and p52 120 and 52 kDa respectively; RELB 70 kDa; loading control Actin 43 kDa. **b**, Blot depicting stabilization of NIK in wild type and all knockout cells treated with vehicle (C) or 10 nM R1881 (T). Representative of two independent experiments. NIK 125 kDa; loading control Actin 43 kDa. **c**, Levels of p100 and p52 protein in DNA and RNA sensors (RIG-I and STING) single and double knockouts in LNCaP cells treated with vehicle (C) or 10 nM R1881 (T). Representative of two independent experiments. NF-kappaB2 p100 and p52 120 and 52 kDa respectively; RELB 70 kDa; loading control Actin 43 kDa. **d**, Heat map depicting fold change for expression of number of genes selected from pan-cancer immune profiling panel. Total RNA extracted from indicated samples was analyzed for 770 immune-related human genes using nCounter human PanCancer Immune Profiling Panel (Nanostring). **e**, Heatmap showing relative expression levels of genes selected from the list of 57 genes which showed higher expression in Nanostring analysis. Total RNA was analyzed using qRT-PCR and data is shown as heatmap from a representative experiment of three independent experiments. **f**, Representative photomicrographs depict CFSE labelled NK-92 cells migrated towards either vehicle (C) or 10 nM R1881 (T) treated LNCaP or LAPC4 cells through 5 µM Boyden chamber. Vehicle control normalized count of NK-92 cells plotted as a mean of four independent measurements on histograms with error bars showing standard deviation. **g**, Immunofluorescence staining of tumor xenograft specimens for LC3B (green) and CXCL10 (red) from tumor bearing mice treated with testosterone cypionate (T) (n=4) in lower panel as compared to tumor isolated from vehicle (C) treated animals (n=4) shown in top panel. **h**, Tumor xenografts specimens from testosterone cypionate treated mice stained with DAPI (grey) and LC3B (green) to show the presence of cytoplasmic autophagosomal DNA. **i**, Representative immunofluorescence images for tumor sections from the vehicle (C) (n=4) or testosterone cypionate treated (T) mice (n=4) stained with Alexafluor488-antiCD57. The number of stained NK cells were counted using Fiji image analysis software and plotted as mean on histogram. **j**. Immunohistochemical staining for CD8 in matched biopsy before and during BAT treatment (n=10). Left photomicrograph shows medium power image of tumor with sparse CD8 cell infiltrate involving tumor, most of the cells seen represent tumor cells. Note increase in extent of infiltrate in treated sample in image on the right. Arrows indicate CD8 positive T cells. Histogram shows area normalized number of CD8 T cells in biopsies collected from PCa patients before and after BAT regimen (n=10). Scale bars: 100 µm (**f**), 10µm (**g**), 2µm (**h**) and 25µm (**i**).

To identify immune genes activated by SupraT and discern those regulated by the STING-TBK1-NFkappaB axis, we performed a Nanostring PanCancer IO 360 Gene Expression analysis using a panel of 770 unique immune-gene signatures. Out of 57 differentially (>2.5 fold) expressed genes, 38 percent of immune genes were induced in a STING-TBK1-NFkappaB dependent manner (Fig. 5d). We validated a panel of chemokines and cytokines that play a role in attracting immune cells using quantitative PCR, and in concurrence to our Nanostring data these genes were altered in a STING-TBK1-NFkappaB dependent manner (Fig. 5e). The gene expression data further indicate that both innate and adaptive immune cells might home to and get activated by SupraT induced cytokines and chemokines. To investigate whether SupraT is able to induce migration of NK cells, we conducted a transwell migration assay with human NK-92 cells and found the SupraT-sensitive cell line significantly increased migration of NK-92 cells upon treatment (Fig. 5f). We next established LNCaP xenografts in athymic nude mice which exhibit robust NK cell activation^29^. In concordance with *in vitro* findings, SupraT led to an increase in autophagosomes, presence of cytoplasmic DNA, and increase in CXCL10 expression (Fig. 5g). The presence of autophagosomal DNA without any autophagy inhibition indicated that the dynamics of autophagosomal degradation differed *in vitro* and *in vivo* (Fig. 5h). Staining for NK cell markers CD57 and CD49b revealed a 17-fold induction in NK cell migration to the tumor bed (Fig. 5i and Extended Data Fig. 9). Prostate tumors are considered as immunologically cold tumors with limited cytotoxic T cell infiltration^30^. The clinical significance of our data was investigated using biopsy from patients undergoing BAT therapy. As seen in Fig. 5j, BAT administration markedly increased infiltration of CD8 T cells. In summary, these results indicate that SupraT may be able to activate immune cells through the NF-kappaB pathway by activating nucleic acid sensors, especially in cells having defects in DNA repair pathways. Our data also indicates that SupraT may be able to induce ferroptosis, a potentially immunogenic cell death mechanism.

## Methods

### Cell culture

LNCaP LAPC4, HEK293T and NK92 cells were purchased from the American Type Culture Collection (ATCC). LNCaP and LAPC4 cells were cultured in phenol red-free RPMI (Thermo Fisher) and HEK293T cells were cultured in DMEM-high glucose (Sigma) supplemented with 10% FBS (Gemini Bio). VCaP cells were cultured in DMEM media (ATCC) containing 1.5 gram/L sodium-bi-carbonate. NK92 cells were cultured in CTS™ AIM V® SFM (Thermo Fisher) with 200 U/mL recombinant IL-2 (Peprotech), 12.5% horse serum (Thermo Fisher), and 12.5% FBS according to manufacturer’s instructions. All cell lines were tested for mycoplasma using the PCR based mycoplasma detection kit (Agilent).

### Knockout/knockdown cell line generation

To generate the gene knockouts, tetracycline induced *cas9* vector (Addgene # 50661) was stably expressed in prostate cancer lines. sgRNAs against target genes were cloned in pLXsgRNA vector (Addgene# 50662) and lentiviral particles for sgRNA were produced in HEK293T cells by co-transfecting pLXsgRNA plasmid with pMD2.G (Addgene# 12259) and psPAX2 (Addgene #12260).(Supplementary table 2a) Viruses were harvested after 48h and cells were infected with lentiviral particles. Transduced cells were treated with 1 µg/mL doxycycline to induce *cas9* before selection with 10 µg/mL blasticidin. Following selection, cells were transferred in 96-well plates to select individual clones. Knockouts were verified by western blots and confirmed clones were expanded and cryopreserved for future experiments.

For NCOA4 knockdown in LNCaP cells, MISSION shRNA constructs (TRCN0000236186 and TRCN0000236187) were purchased from Sigma and lentiviral partilces were generated in HEK293T cells by contransfecting shRNA plasmid construct with pMD2.G (Addgene#12259) and psPAX2 (Addgene#12260). LNCaP cells were infected with NCOA4 lentivirus and were selected with 1.0 µg/mL puromycin. Level of NCOA4 knockdown was measured by western blots and confirmed cells were expanded and cryopreserved and used for the experiments.

### Quantitative real-time PCR

Total RNA was extracted from cells using Trizol reagent (Thermo Fisher) as per manufacturer’s instructions. cDNA was synthesized from RNA by reverse transcription using SuperScript IV reverse (Thermo Fisher). Quantitative PCR was performed on samples mixed with SYBR master mix (BioRad) and gene-specific primers using a real-time thermocycler (Biorad). Data analysis was performed using the ΔΔCt method, and fold change (2^^-^ΔΔ^Ct^) was calculated after double normalization with housekeeper gene and respective untreated control. Each PCR reaction were run in triplicates. Details of primer sequences used in qPCR are provided in a supplementary table 2b.

### Immunoprecipitation

NCOA4-FLAG-HA (CTAP) plasmid was received as a gift from Prof. J Wade Harper (Harvard Medical School, Boston). LNCaP cells were transfected with NCOA4-FLAG plasmid for immunoprecipitation studies or kept as untransfected control (C). NCOA4-FLAG transfected cells were either treated with 10 nM R1881 (T) or kept as untreated control (C*). Following treatment, cells were washed with 3 mL PBS, harvested in ice-cold PBS, and pelleted using a refrigerated centrifuge. Cells were lysed with 750 μL 1X lysis buffer (Promega) containing protease and phosphatase inhibitor for 20 minutes on ice. After lysis, the lysates were centrifuged at 13200 ⨯ g at 4 °C, and 10 percent of supernatant was stored as input. Remaining supernatant was used for immunoprecipitation using anti-FLAG M2 agarose beads (Thermo Fisher) as per manufacturer’s instructions. Affinity separated FLAG-tagged protein was eluted using 35 μL IgG elution buffer (Thermo Fisher), denatured with equal volume of Laemlli buffer, and stored at −80 °C before western blot analysis.

### Western blotting

Cell lysates were prepared either in 1X denaturing lysis buffer (Cell Signaling #9803) or non-denaturing lysis buffer (50 mM HEPES pH 7.4, 150 mM NaCl, 5 mM EDTA, 10% glycerol, 1% Nonidet P-40, and supplemented with phosphatase and protease inhibitors). For denaturing gels, lysates were mixed with Laemelli buffer containing beta mecaptoethanol, boiled for 5mins and stored at −80 °C until use. Native lysates were mixed with native loading buffer (BioRad) and semi-native lysates were mixed with Laemelli buffer without betamercaptoethanol and stored in −80 °C without boilng. Native gels were resolved as described^31^; denaturing and semi-native gels were resolved on 4-15% precast polyacrylamide gels (BioRad) followed by transfer to PVDF membrane. Membranes were blocked in 5% nonfat milk in wash buffer (Tris-buffered saline+0.1% Tween-20). After blocking, immunoblotting was performed with primary antibodies overnight at 4 °C followed by incubation with HRP-conjugated secondary antibodies for 1h at room temperature. Blots were analyzed using chemiluminescence method. Blots were developed using ECL Western blot detection reagent (GE Healthcare) for highly expressed proteins and SuperSignal West Femto (Thermo Fisher) reagent for low expression proteins. The antibodies used are listed in a supplementary table 3.

### Immunofluorescence

Cells were grown on sterile coverslips in low density and were treated with R1881 (T) or vehicle control (C). After treatment, media was removed and cells were fixed and permeabilized using chilled methanol for 10 min in deep-freezer followed by 10 min fixation in 10% natural buffered formalin at room temperature. Post-fixation, cells were blocked overnight in sterile-filtered 5% BSA in PBS at 4°C. Cells were incubated with primary antibody, followed by suitable fluorochrome tagged secondary antibody at 4 °C for overnight. Both, primary and secondary antibodies were diluted with 5% BSA in PBS. Cells were postfixed with 10 % NBF for 2-3 min before mounting on a clean glass slide using Vectashield antifade mounting media (Vector Laboratories). Image acquisition was performed on the LSM 700 laser confocal microscope (Zeiss). Images were analyzed using the image processing package Fiji.

### Measurement of cell death

To determine cell death, cells were stained with 0.4% trypan blue (Sigma)and incubated for 2-3 min at room temperature to allow dead cells to take up the blue stain. Cell viability was measured using an automated cell counter, Cellometer, (Nexcelom). Dead cells were calculated from subtracting the percentage of viable cells from 100. Percentage of dead cells were plotted on histograms.

### NK cell migration assay

This assay was performed in a transwell co-culture set up, using tumor cells in the bottom chamber (culture plate) and NK-92 cells in the upper chamber (insert) (Corning). LNCaP and LAPC4 cells were first seeded in a 24-well plate for 24h (day 1). The following day, tumor cells were treated with vehicle (C) or 10 nM R1881 (T) for 72h. After 72h, NK-92 cells, stained with 5 μM CFSE (Biolegend) for 10 min at 37 °C, were then added to a 5-micron pore size insert placed in each culture well already containing the vehicle/R1881 treated tumor cells. The culture plate was incubated for 6h and microscopy was performed to image the CFSE labelled NK-92 cells that migrated to the lower chamber of the plate. Images were analyzed using the Image J software to determine the cell count.

### Cell cycle analysis

Harvested cell pellets were resuspended in ice-cold 70% ethanol and incubated at −20 °C overnight. Cells were then washed with 1X PBS, resuspended in 50 μg/ml propidium iodide (Sigma) in PBS with 50 μg/ml RNase A (Sigma), and incubated for 30 min at room temperature. Cell cycle analysis was performed using BD FACSCelesta cytometer equipped with BD FACS DIVA. Data were analyzed using the FlowJo software version 10 (FlowJo). Distribution of cells in the cell cycle phases was calculated using the Watson-Pragmatic algorithm.

### NanoString immune profiling assay

Total RNA was isolated from vehicle/R1881 treated cells using Trizol (Thermo Fisher), and RNA quality was determined using the Bioanalyzer (Agilent). Gene signature was determined by using the NanoString PanCancer Immune Profiling Panel according to the manufacturer’s instructions (NanoString Technologies). House-keeping genes and built-in positive controls were used for data normalization and quality control assesment. Data analysis and gene expression analysis was performed using the nSolver 4.0 software (NanoString Technologies).

### Luciferase reporter assay

Dual-Luciferase® reporter assay (Promega) was used to measure NF-kB, STING, and AIM-2 promoter activity. LNCaP and LAPC4 cells were transfected with Ig-IFN-Luc and Renila-Luc plasmids in 12-well plates and were incubated with vehicle or 10 nM R1881 treatment for 72h. Post-treatment, cells were washed with PBS, followed by lysis with 250 μL 1X Passive lysis buffer (Promega) at room temperature for 15 min. The lysate was cleared by isolating cell debris by centrifugation and stored at −80 °C unless used immediately for luminescence measurement. For luminometric measurement, 100 μL LARII buffer containing D-luciferin was dispensed in each well of the luminometer plate followed by 20 μL of cell lysate to measure *Firefly* luminescence. After measuring firefly luminescence, 100 μL Stop & Glo ® reagent containing quencher and substrate for *Renilla* luciferase was added to measure respective responsive luminescence. Data was normalized using individual well’s *Renilla* luminescence.

### Bioplex assay for chemokine and cytokine measurement

Concentration of 17 chemokines, cytokines, and growth factors in culture supernatants of control or R1881 treated LNCaP cells was determined using the Bioplex Pro Human Cytokine 17-plex assay system (Bio-Rad) according to manufacturer’s protocol. Culture supernatant from vehicle and R1881 treated LNCaP and LAPC4 cells was collected at 3 and 6 day timepoints, and stored at −80 °C until further analysis. Data was collected on the Bio-Rad BioPlex 200 instrument and analyzed using the Bio-Plex Manager (Bio-Rad Laboratory).

### Autophagosome Flux measurement

Cells were plated on 60 mm dishes to obtain 80% confluence on the next day. Cells were transfected with tandem mRFP-GFP fluorescent-tagged LC3 (ptfLC3) (Addgene #21074). 24h post-transfection, cells were trypsinized and plated on sterile glass bottom dishes at a 40–50% confluency and treated with androgens. Cells were then imaged using a live cell Zeiss LSM780-FCS Single-point, laser scanning confocal microscope. Images were processed using the ImageJ software. Cells with predominantly yellow RFP/GFP (autophagosome) or red RFP (autolysosome) punctae were counted and analyzed.

### Mouse xenografts and treatment

Using a Johns Hopkins Animal Care and Use Committee approved protocol, adult athymic nude mice were inoculated subcutaneously in the flank with the LNCaP human prostate cancer cell lines in 200 μL of Matrigel. Mice were divided into two groups and the treatment group was implanted with 2 one cm long silastic implants filled with testosterone as described previously^6^. Tumors were harvested 2 and 4 days post-treatment and fixed in 10% buffered formalin and processed for IHC and H&E staining. NK cells were identified using antiCD57-PE (Santa Cruz Biotechnology) and antiCD49b-FITC (Santa Cruz Biotechnology) antibodies. Stained sections were imaged using Zeiss LSM 700 laser confocal microscope. Images were analyzed using the image processing package Fiji. NK cells were counted per field and plotted as mean values.

### Patient Materials

Biopsies from metastatic lesions were obtained from patients with castration-resistant prostate cancer, under an IRB-approved prospective protocol (clinicaltrials.gov: NCT03554317) examining the use of SupraT as a treatment for metastatic castration-resistant prostate cancer (mCRPC). These patients had previously been treated with standard androgen deprivation therapy as well as next-generation anti-androgen therapy using abiraterone and/or enzalutamide. Biopsies were typically obtained from soft-tissue metastatic sites (lymph nodes, liver, and lung) using a 18-gauge core biopsy needle, and were collected before and 12 weeks after starting the SupraT treatment. For each metastatic biopsy, at least 2 fresh cores were collected for immediate flash-freezing in liquid nitrogen (Frozen tissue method), and at least 2 separate cores for formalin-fixed and paraffin-embedded sections (FFPE method). To minimize tissue autolysis, a dedicated tissue technician was called to the biopsy suite to initiate processing steps, including FFPE preparation within 30 minutes of biopsy collection. All samples were evaluated for adequacy and assessment of tumor-to-normal ratio by an expert urological pathologist. FFPE specimens were processed according to the standard procedures at the Johns Hopkins pathology department.

### Immunohistochemical Staining and CD8 density measurement by image analysis

Chromogenic IHC for CD8 was performed as follows. Sections of 4 micron thickness were cut and placed on super frost plus slides. Paraffin Sections were baked on a hot plate at 60 °C for 10 minutes, dewaxed using xylene, rehydrated in a series of graded alcohols to distilled water, and finally rinsed in distilled water with 0.1% Tween 20. Slides were transferred to a glass jar filled with suitable antigen retrieval solution. The glass jar was irradiated in a microwave oven at full power for 1 minute, followed by 15 minutes at power level 20. Slides were cooled for 5 minutes at room temperature and washed 2X in tris-buffered saline with 0.1% Tween 20 (TBST). Tissues were subjected to endogenous peroxidase blocking using hydrogen peroxide for 5 minutes. Slides were then incubated with CD8 (DAKO, Clone: C8/144B) antibody for 45 minutes at room temperature, rinsed with TBST, and incubated with the secondary antibody (PowerVision Poly-HRP Anti-Mouse IgG, Leica, PV6119) for 30 minutes. Following incubation with the chromogen, 3-Amino-9-Ethylcarbazole (AEC) for 20 minutes, slides were counterstained with hematoxylin. After counterstaining, slides were washed with tap water for 2 minutes and distilled water for 1 minute and mounted using VectaMount AQ, Vector H-5501.

Whole biopsy slides stained for CD8 were scanned on a Roche-Ventana DP200 whole slide scanner and analyzed using the HALO 3.0 (Indica Labs) software. Regions of interest (ROI) consisted of tumor tissue that was delineated manually by a pathologist with expertise in prostate pathology. In cases with clear lymph node tissue from lymph node biopsies, regions were chosen to avoid encompassing the lymphoid tissue apart from the tumor. T cells were delineated using the cytonuclear IHC module in HALO. CD8 density was calculated as the number of T cells per mm^2^ of ROI. Cell density measured using HALO were verified by manual counting in a subset of cases.

### Statistical analysis

Suitable central tendency values were calculated for all the quantitatively measurable variables and were used for the analysis of statistical significance. Parametric analysis was performed to compare mean values after estimating the normal distribution, unless otherwise specified. These analyses were performed using the Prism version 6.0 (GraphPad Software) and a value of *p≤0*.*05* was considered significant.

---

**Extended Data Fig. 1:**
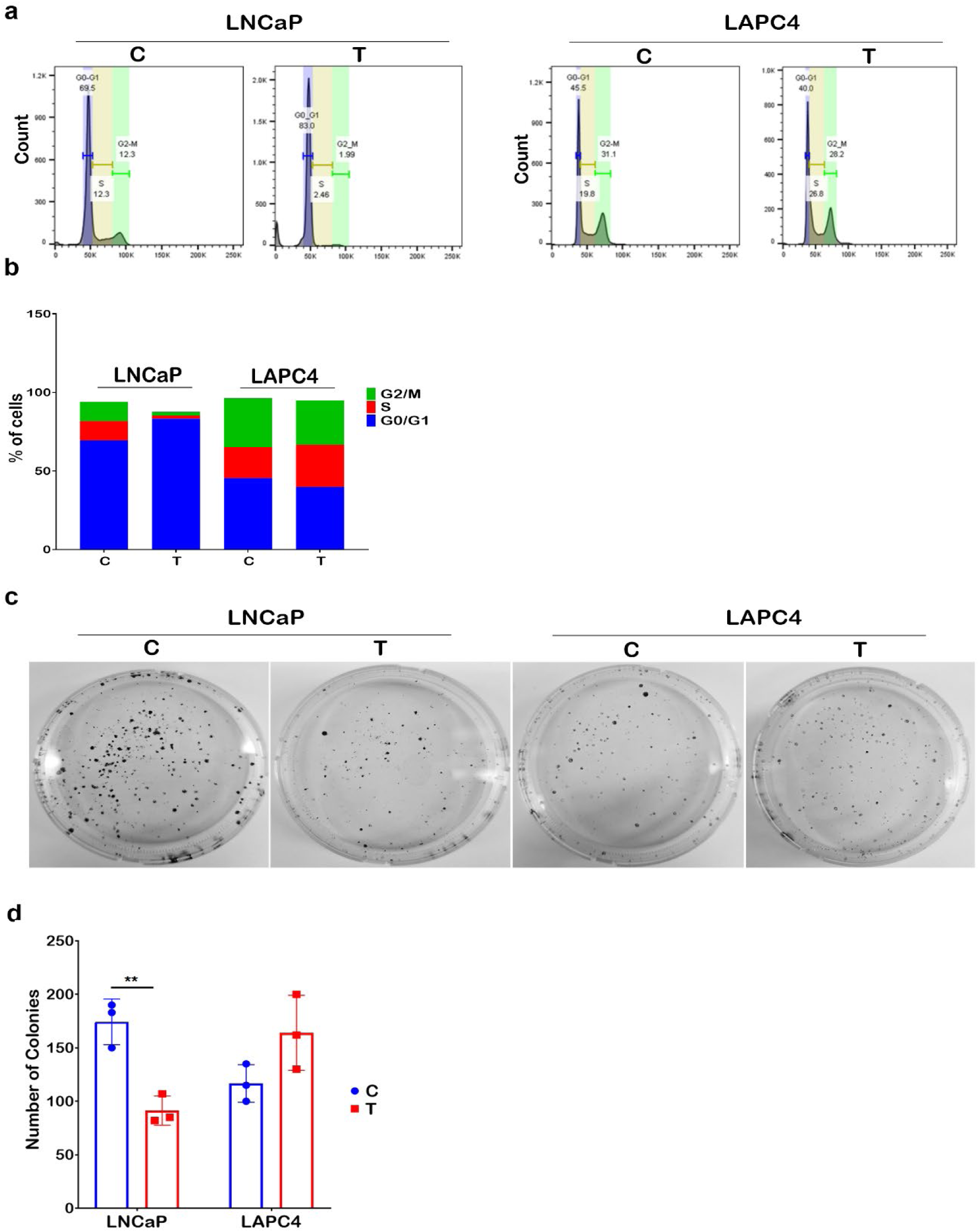
Effect of SupraT on cell cycle behavior and clonogenic potential in PCa cells. Histogram plot showing **a**, cell cycle distribution for LNCaP and LAPC4 cells treated with vehicle (C) or 10 nM R1881 (T) for 72 h. **b**, Bar graph showing percentage of cells in different cell cycle phases from a representative experiment for LNCaP and LAPC4 cells from two independent experiments. **c**, Representative images of crystal violet stained cell colonies cultured for 14 days from three independent experiments. Images were analyzed using Colony Counter plugin of the image analysis software, Fiji. **d**, Bar graph showing an average number of colonies in vehicle control (C) or 10 nM R1881 treated (T) LNCaP and LAPC4 cells for 72h and subsequently plated for 14 days.

**Extended Data Fig. 2:**
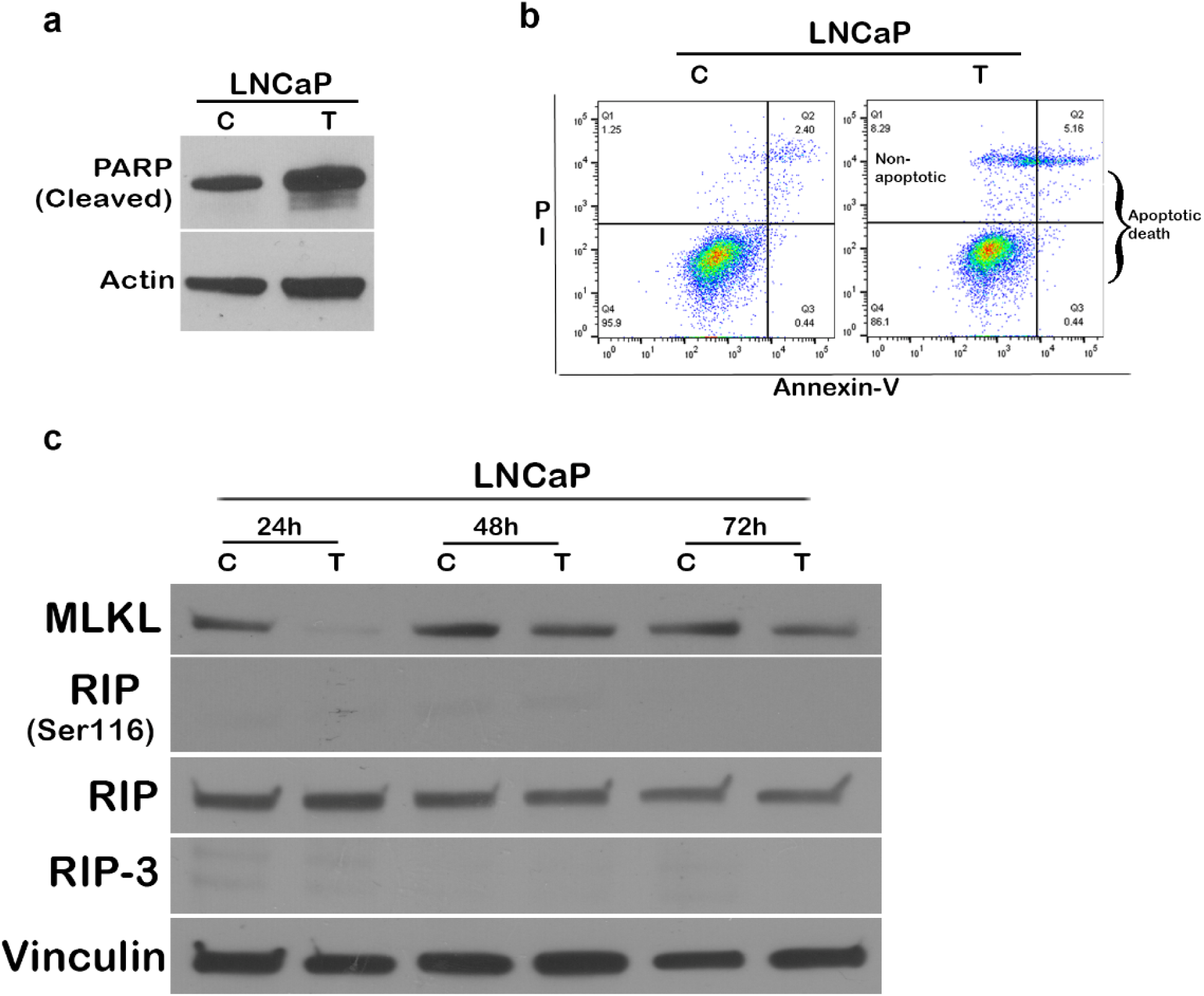
Measurement of apoptotic and necrotic cell death in SupraT treated LNCaP cells. Representative western blot image from two independent experiments for cleaved PARP in **a**, LNCaP cells treated with vehicle (C) or 10 nM R1881 (T) for 72h. Cleaved PARP 89 kDa; loading control Actin 43 kDa. **b**, Annexin-PI analysis of LNCaP cells treated with vehicle (C) or 10 nM R1881 (T) for 48h. Representative flowcytogram from two independent experiments. Annexin positive cells indicates cells dying through apoptosis and PI alone positive cells indicate cells dying through non-apoptotic mechanism. **c**, Western blot analysis for necrotic cell death markers MLKL, RIP, RIP (Ser116) and RIP3 in vehicle (C) or 10 nM R1881 treated (T) LNCaP cells for 24h, 48h, and 72h time points. Representative of two independent experiments. MLKL 54 kDa; Phospho-RIP 78-82 kDa; RIP 78 kDa; RIP3 46-62 kDa; loading control Vinculin 124 kDa.

**Extended Data Fig. 3:**
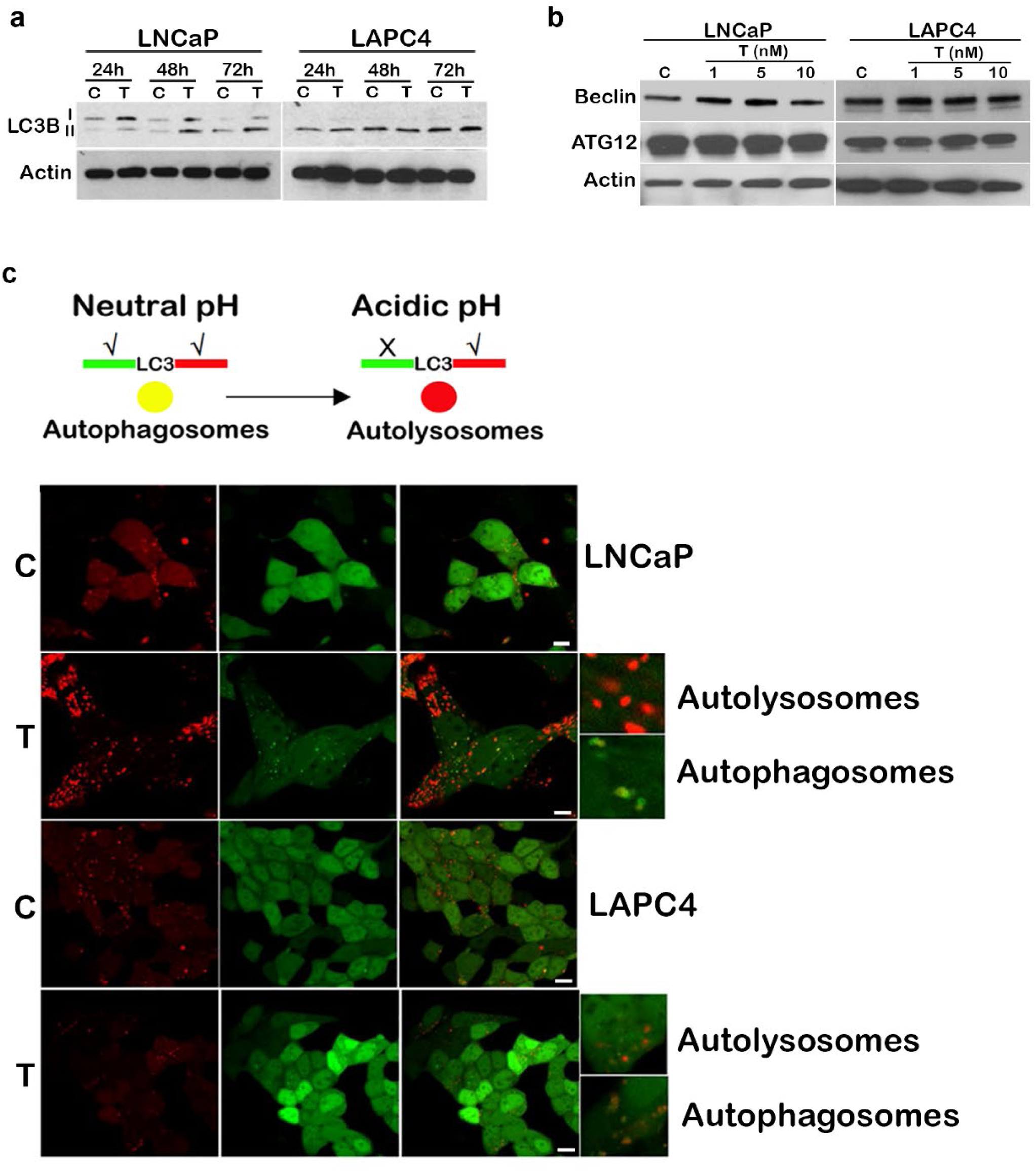
Measurement of autophagy in SupraT treated PCa cells. **a**, Immunoblot for LC3B-II in LNCaP and LAPC4 cells treated with vehicle (C) or 10 nM R1881 (T) for 24h, 48h, and 72h. Representative of two independent experiments. LC3B 14,16 kDa; loading control Actin 43 kDa. **b**, immunoblot for measurement of global autophagy through Beclin and ATG12 in cells treated with vehicle (C) or R1881 (1, 5, and 10 nM) (T). Representative of two independent experiments. Beclin 60 kDa; ATG 12 55 kDa; loading control Actin 43 kDa. **c**, Autophagy flux sensor RFP-LC3-GFP in LNCaP and LAPC4 cells treated with either vehicle (C) or with 10 nM of R1881 (T) for 72h and imaged using confocal microscopy to visualize autolysosomes (red) and autophagosomes (red and green double-positive) puncta. A representative single confocal section is shown from three independent experiments. Scale bars: 10 µm (**c**).

**Extended Data Fig. 4:**
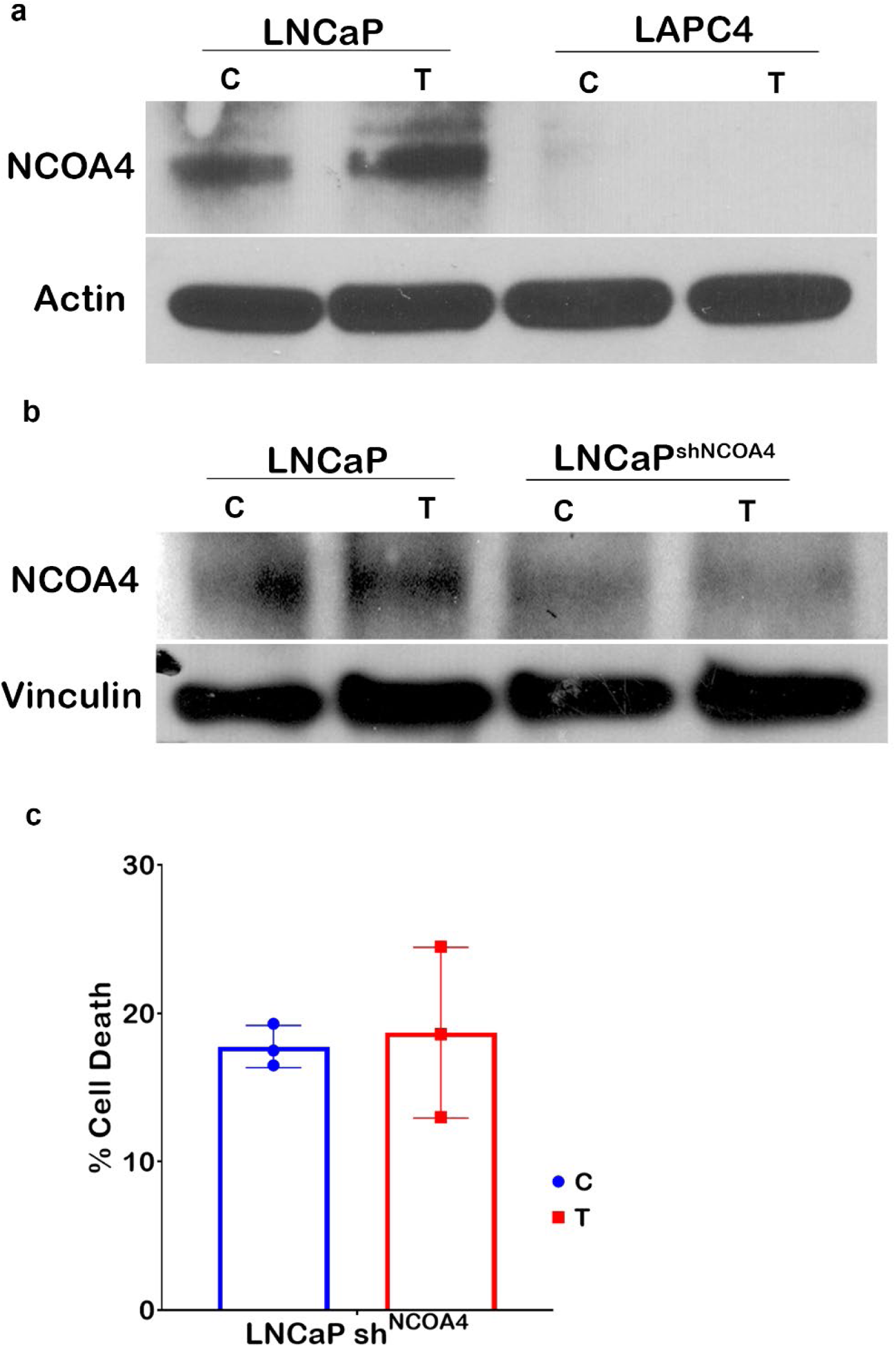
NCOA4 is required for SupraT induced cell death in LNCaP cells. **a**, Western blot-based measurement of NCOA4 protein in vehicle (C) or 10 nM R1881 treated (T) LNCaP and LAPC4 cells for 72h. Representative of two independent experiments. NCOA4 70 kDa; loading control Actin 43 kDa. **b**, Western blot analysis of NCOA4 in control and NCOA4 knockdown LNCaP cells after vehicle (C) or 10 nM R1881 treatment (T). Representative of two independent experiments. NCOA4 70 kDa; loading control Vinculin 124 kDa. c, Trypan blue based viability measurement for vehicle (C) or 10 nM R1881 treated (T) NCOA4 knockdown LNCaP cells. Histogram bar shows mean % dead cells and error bar indicates standard deviation from three measurements.

**Extended Data Fig. 5:**
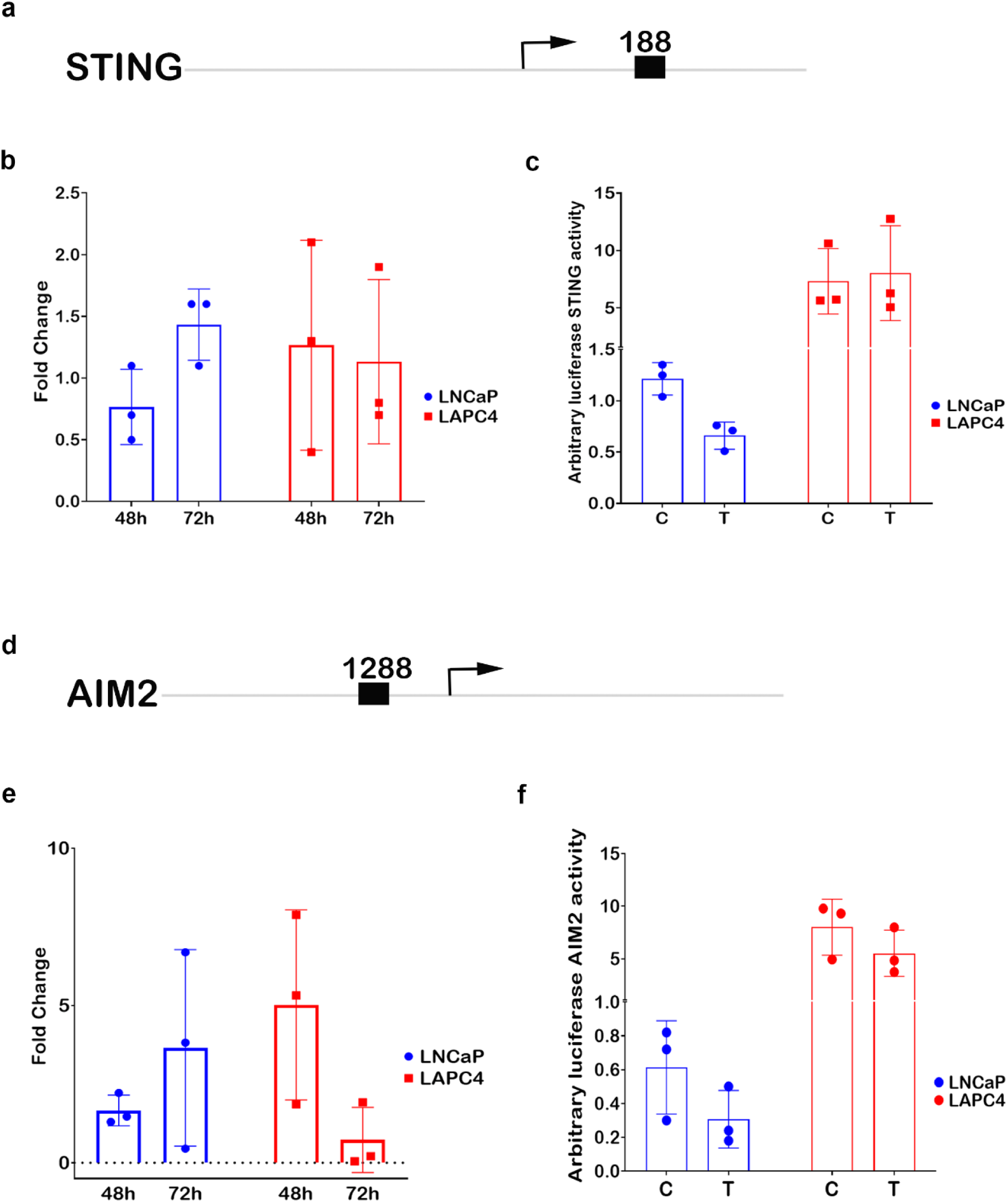
Regulation of STING and AIM2 by SupraT treatment. **a**, Presence of androgen binding motifs at STING promoter sequence. Arrow indicates transcription start site and numbers indicate base pairs away from the transcription start site. **b**, Level of STING transcripts by quantitative RT-PCR in LNCaP and LAPC4 cells treated with 10 nM R1881 for 48, or 72h. Histogram bars indicate mean fold change calculated using vehicle-treated control cells. **c**, STING promoter activity by luciferase-based measurement in vehicle control (C) and 10 nM R1881 treated (T) LNCaP and LAPC4 cells after 72 h. Histogram indicates mean arbitrary firefly luminescence indicative of STING promoter activity from three independent measurements with error bars showing standard deviation. **d)** Presence of androgen binding motif at AIM-2 promoter sequence. Arrow indicates transcription start site and numbers indicate base pairs away from the transcription start site. **e**, Level of AIM2 transcripts by quantitative RT-PCR in LNCaP and LAPC4 cells treated with 10 nM R1881 for 48, and 72h. Histogram bars indicate the mean of fold change calculated using vehicle-treated control cells. **f**, AIM2 promoter activity by luciferase-based measurement in vehicle control (C) and 10 nM R1881 treated (T) LNCaP and LAPC4 cells after 72 h. Histogram indicates mean arbitrary firefly luminescence indicative of AIM2 promoter activity from three independent measurements with error bars showing standard deviation. Double reporter assay was utilized for measurement and Renilla luminescence values were used for normalization.

**Extended Data Fig. 6:**
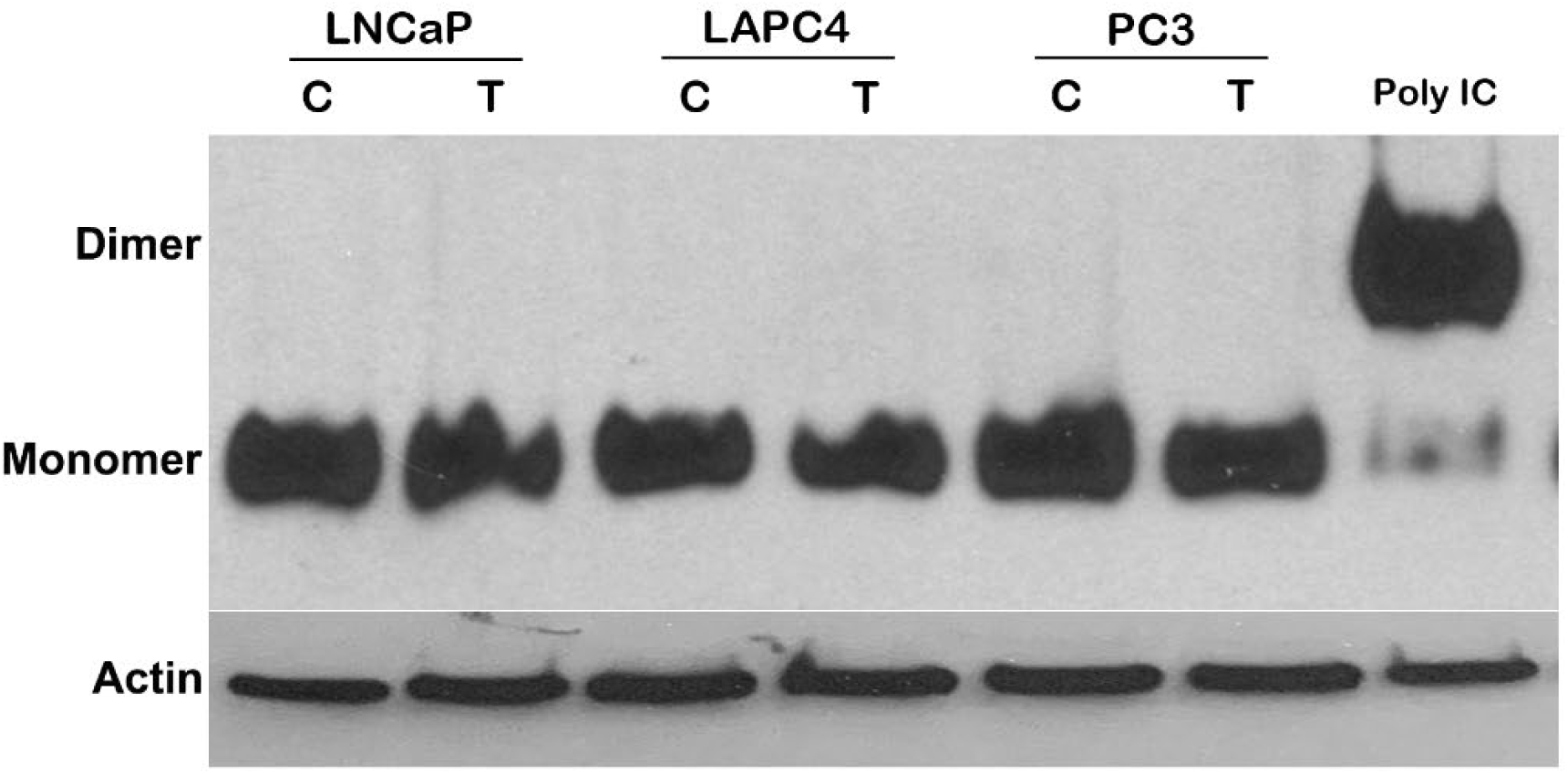
Activation of IRF3 in PCa cells by SupraT. Native western blot analysis of PCa cells for IRF3 dimerization in vehicle-treated control (C) or 10 nM R1881 treated (T) LNCaP, LAPC4, or AR negative PC3 cells. Poly: IC treated cells were used as a positive control for the IRF3 dimerization. Representative of two independent experiments; lysate form Polyinosinic-polycytidylic acid treated PC3 cells was used as positive control; loading control Actin 43 kDa.

**Extended Data Fig. 7:**
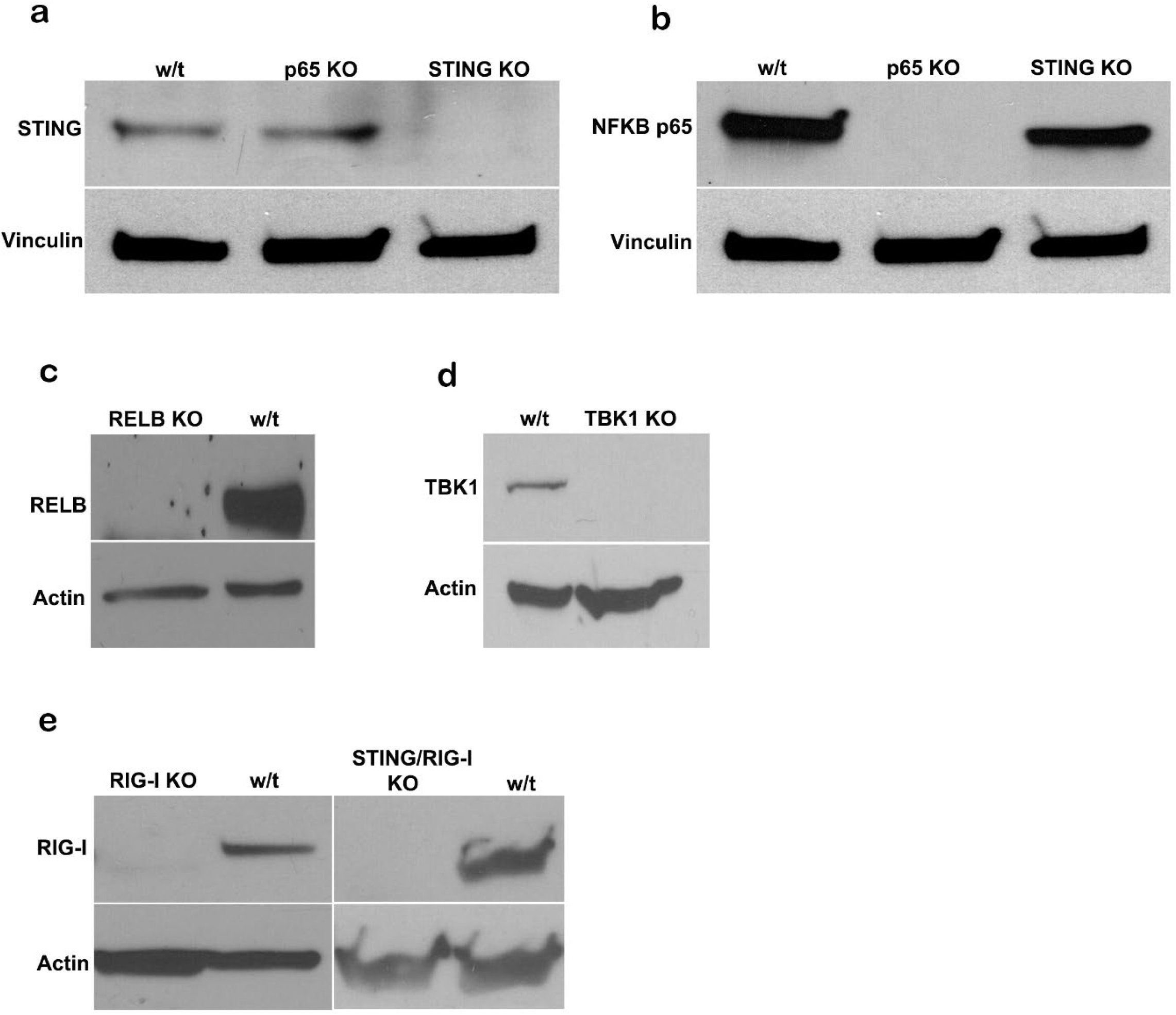
Western blot confirmation for all the knockouts generated through CRISPR-Cas based approach. Protein lysates were resolved through SDS-PAGE, and immobilized protein(s) was probed with respective antibody to screen k7nockouts for **a**, STING, **b**, NF-kappaB -p65 **c**, RELB, **d**, TBK1, and **e**, RIG-I in LNCaP, as well as LNCaP with STING knockout cells. STING 33-35 kDa; NF-kappaB p65 65 kDa, RELB 70 kDa, TBK1 84 kDa, RIG-I 102 kDa; loading control Vinculin 124 kDa and Actin 43 kDa.

**Extended Data Fig. 8:**
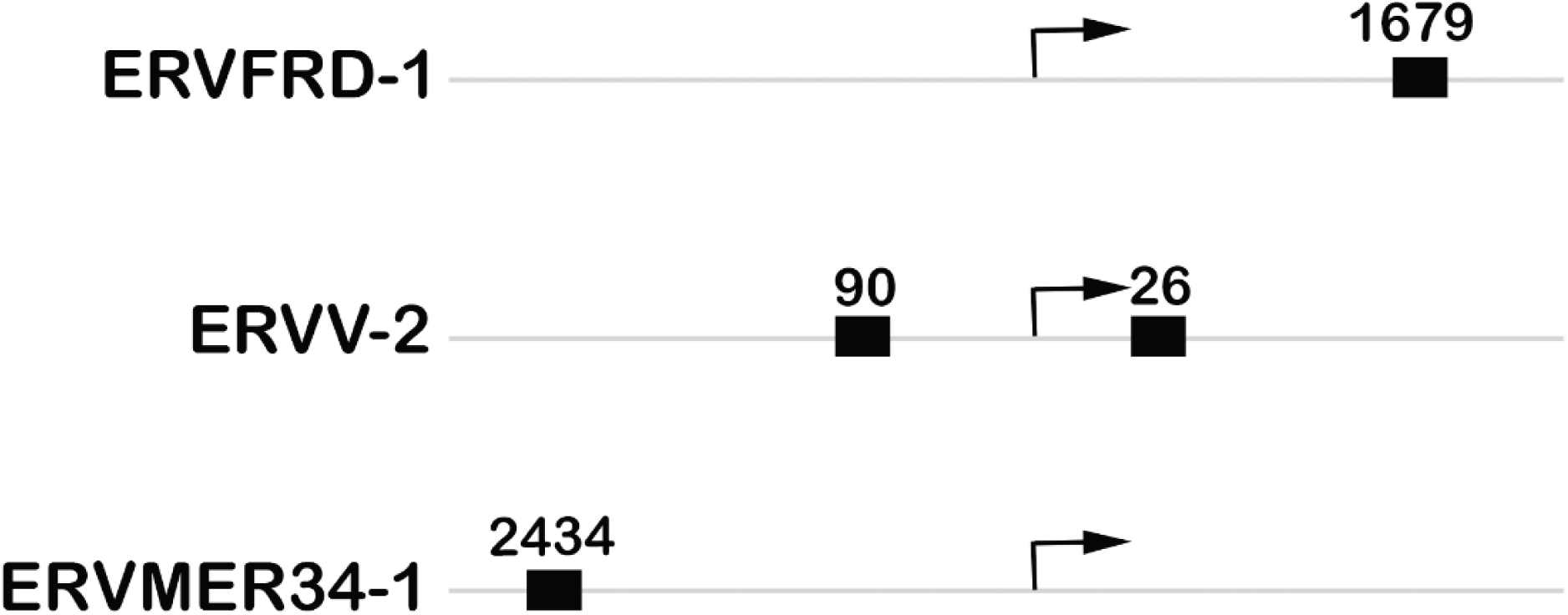
Androgen receptor response elements in endogenous retroviruses. Genomic regions of endogenous retroviruses depict the presence of androgen response elements close to the transcription start site (depicted by arrow). Numbers indicate base pairs away from the transcription start site.

**Extended Data Fig. 9:**
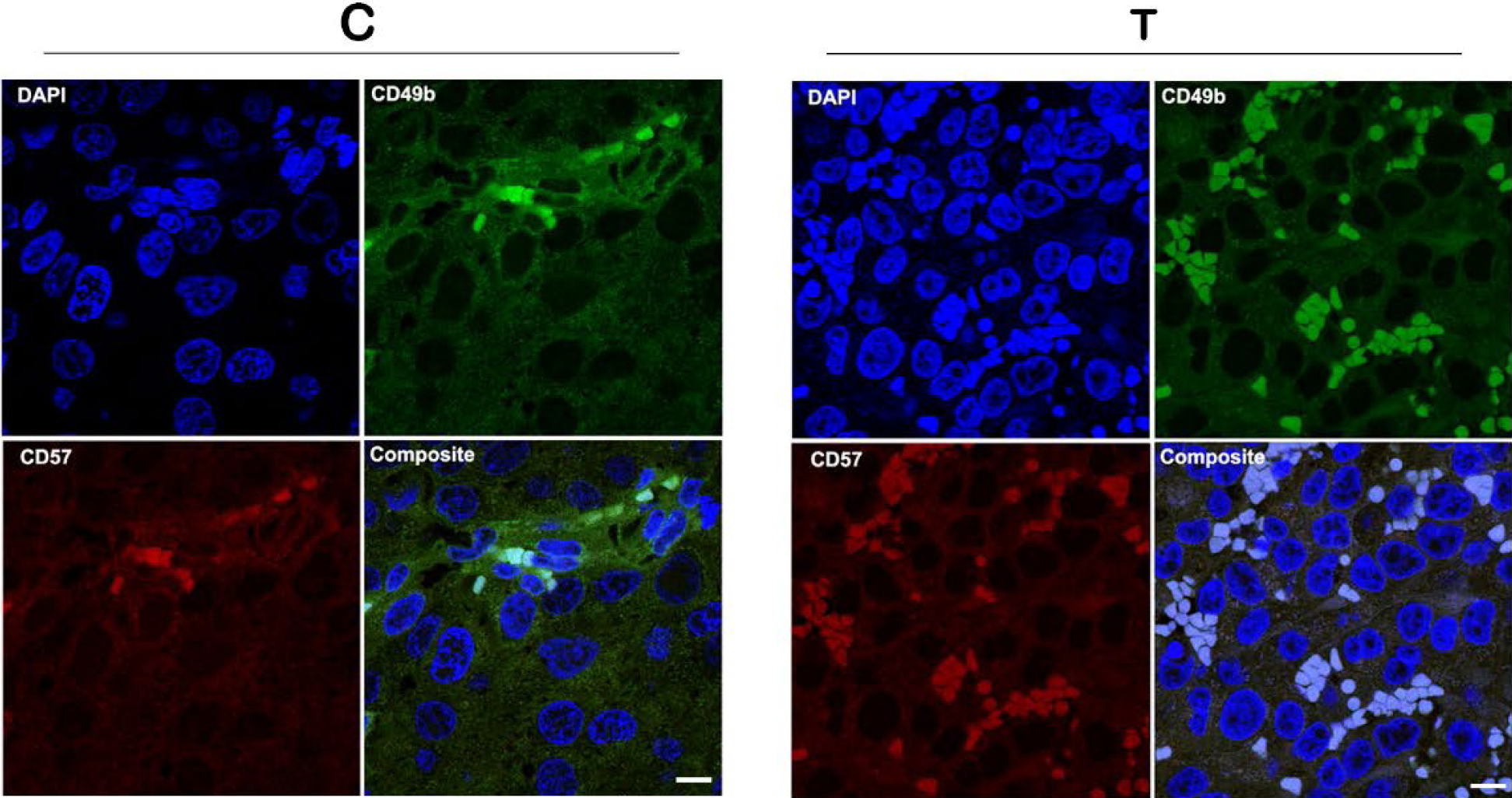
NK cell surface markers show activated NK cell in tumor xenografts. Representative immunofluorescence images for tumor sections from the vehicle (n=4) or testosterone-treated mice (n=4) stained with PE-antiCD57 and FITC-antiCD49b. Scale bars: 10 µm.

**Extended Data Table 1:** DNA repair gene mutation analysis. Table showing results from the search of Cosmic cell line database for mutations in DNA repair genes in PCa cell lines LNCaP, VCaP, and LAPC4.

|  | *BRCA2* | *ATM* | *MSH2* | *MSH6* | *FANCA* |
| --- | --- | --- | --- | --- | --- |
| **LNCaP** | p.D946fs*14 | p.Q912* | Homologous<br>Deletion |  |  |
| **VCaP** |  |  |  | p.F1088fs*5 | Homologous<br>deletion |
| **LAPC4** |  |  |  |  |  |

## Acknowledgement

We thank Dr. Sabatini and Dr. Lander for pCW-Cas9 (Addgene#50661) and pLXsgRNA (Addgene#50662), Dr. Trono for pMD2.G (Addgene 12259) and psPAX2 (Addgene#12260), and Dr. Wade Harper for the NCOA4α-pHAGE-C-FLAG-HA plasmid. This work is supported by the DOD grants W81XWH1910724 to SK and W81XWH-14-2-0189 to SD; Allegheny Health Network-Johns Hopkins Cancer Research Fund, Shared Instrumentation grant S10OD016374, and the NCI CORE Grant P30CA006973. ESA and SK are partially supported by a PCF 2018 Challenge Award.

## Ethics Declaration

ESA is a paid consultant/advisor to Janssen, Astellas, Sanofi, Dendreon, Pfizer, Amgen, Eli Lilly, Bayer, AstraZeneca, Bristol-Myers Squibb, Clovis, and Merck; and has received research funding from Janssen, Johnson & Johnson, Sanofi, Dendreon, Genentech, Novartis, Tokai, Bristol Myers-Squibb, AstraZeneca, Clovis, and Merck.

