## Supplementary table 1 for "Supraphysiological testosterone induces ferroptosis and activates NF-kappaB mediated immune pathways in prostate cancer through nucleophagy"

Supplementary Table: 1 Bioplex assays measurement of chemokines

| Treatment | Time | LNCaP (pg/mL) | | | LAPC4 (pg/mL) | | |
| --- | --- | --- | --- | --- | --- | --- | --- |
|  |  | CXCL10 | CCL2 | CCL5 | CXCL10 | CCL2 | CCL5 |
| C | d3 | 6.5 | BDL | BDL | BDL | BDL | BDL |
| T |  | 23.2 | 6.0 | BDL | BDL | BDL | BDL |
| C | d6 | 44.5 | 14.6 | 10.0 | BDL | BDL | BDL |
| T |  | 116.8 | 31.9 | 20.4 | BDL | BDL | BDL |

C = Vehicle control

T = 10 nM R1881

BDL= Below detection limit
