## Supplementary table 2 for "Supraphysiological testosterone induces ferroptosis and activates NF-kappaB mediated immune pathways in prostate cancer through nucleophagy"

| Supplementary table 2a: sgRNA sequences | | |
| --- | --- | --- |
|  |  | 5'-3' sequence for sgRNA |
| pLXsgXho1 | F1 | AAACTCGAGTGTACAAAAAAGCAGGCTTTAAAG |
| pLXsgNhe1 | R2 | AAAGCTAGCTAATGCCAACTTTGTACAAGAAAGCTG |
|  |  | sgRNA1 Sequences |
| *STING_sgRNA1* | R1 | CCAGTAGCTGCCCCGGTACCGGTGTTTCGTCCTTTCC |
|  | F2 | GGTACCGGGGCAGCTACTGGGTTTTAGAGCTAGAAATAGCAA |
| *STING-sgRNA2* | R1 | GGCAGGCACTCAGCAGAACCGGTGTTTCGTCCTTTCC |
|  | F2 | GGTTCTGCTGAGTGCCTGCCGTTTTAGAGCTAGAAATAGCAA |
| *p65_sgRNA1* | R1 | CCTGCCGCCTGTCCTTTCTCGGTGTTTCGTCCTTTCC |
|  | F2 | GAGAAAGGACAGGCGGCAGGGTTTTAGAGCTAGAAATAGCAA |
| *p65_sgRNA2* | R1 | GCTCCAGGTCCCGCTTCTTCGGTGTTTCGTCCTTTCC |
|  | F2 | GAAGAAGCGGGACCTGGAGCGTTTTAGACGTAGAAATAGCAA |
| *RELB_sgRNA1* | R1 | CGGCTCAGGTGGGTCCCAGCGGTGTTTCGTCCTTTCC |
|  | F2 | GCTGGGACCCACCTGAGCCGGTTTTAGAGCTAGAAATAGCAA |
| *RELB_sgRNA2* | R1 | TCCGGCGCATCTGTCCCTGCGGTGTTTCGTCCTTTCC |
|  | F2 | GCAGGGACAGATGCGCCGGAGTTTTAGAGCTAGAAATAGCAA |
| *TBK1_sgRNA1* | R1 | CTTCCAGGTTGATCTTCATCGGTGTTTCGTCCTTTCC |
|  | F2 | GATGAAGATCAACCTGGAAGGTTTTAGAGCTAGAAATAGCAA |
| *TBK1_sgRNA2* | R1 | TTTTCTGCTTTCTGTACTCCGGTGTTTCGTCCTTTCC |
|  | F2 | GGAGTACAGAAAGCAGAAAAGTTTTAGAGCTAGAAATAGCAA |
| *RIG-I_sgRNA1* | R1 | CAGTTCACTGAACTTGTTCCGGTGTTTCGTCCTTTCC |
|  | F2 | GGAACAAGTTCAGTGAACTGGTTTTAGAGCTAGAAATAGCAA |
| *RIG-I_sgRNA2* | R1 | CCTCAGTTGCTGATGAAGGCGGTGTTTCGTCCTTTCC |
|  | F2 | GCCTTCATCAGCAACTGAGGGTTTTAGAGCTAGAAATAGCAA |
| Supplementary table 2b: qPCR primers | | |
| Gene |  | 5'-3' sequence |
| *SLC7A11* | F' | GTGGGGTCCTGTCACTATTT |
|  | R' | TGCAGGGCGTATTATGAGGA |
| *GPX4* | F' | TGTAACCAGTTCGGGAAGCA |
|  | R' | TGATGGCATTTCCCAGGATG |
| *ALOX5* | F' | ACACCGACGTAAAGAACTGG |
|  | R' | ACAGGTTCTCCATCGCTTTG |
| *ALOX15* | F' | TCTTCTGAGGGGACACTTGA |
|  | R' | CAGTGCTCATTATCTGGTCG |
| *ACSL3* | F' | CAATTTCCGAAGTGTGGGAC |
|  | R' | CACTTTGGCCCCCAATAAGA |
| *ACSL4* | F' | TCTCCTCAGACACACCGATT |
|  | R' | GTATAACCGCCTTCTTGCCA |
| *NCOA4* | F' | CGAGAGGTCAAAGCTCAGAT |
|  | R' | ACTGGCCCAATAACGAGTAG |
| *GLS2* | F' | GGGCAGAAATTCGGAACAAG |
|  | R' | TTCACTTTGCAAGCCTCGAT |
| *DPP4* | F' | TCGGCGCTCACTAATGTTTA |
|  | R' | AACCTTCCACGGTGTCTTCA |
| *CHAC1* | F' | CTCCTTGAAGATCATGAGGGCT |
|  | R' | AAGTATTCAAGGTTGTGGCCG |
| *PTGS2* | F' | GCAGGCTAATACTGATAGGAGAG |
|  | R' | ATGCCAGTGATAGAGGGTGT |
| *CXCL11* | F' | TGGTTACGGTGGAGACATTG |
|  | R' | TGCCAGTATCCCATAGCGT |
| *IL32* | F' | TCCCGAAGGTCCTCTCTGAT |
|  | R' | GGCACCGTAATCCATCTCTT |
| *TNFRSF9(CD147)* | F' | GACACTCTCCGCAGATCAT |
|  | R' | CTTCTTCTGGAAATCGGCAG |
| *CCL2* | F' | CACTCCACAACCCAAGAATC |
|  | R' | AGGGGTAGAACTGTGGTTCA |
| *CD68* | F' | TTGGGCAAAGTTTCTCCTGC |
|  | R' | CGTGTCGAGGAAATAAGGGT |
| *IL8* | F' | GGGCCAAGAGAATATCCGAAC |
|  | R' | ACAAGGACTTGTGGATCCTG |
| *CCL8* | F' | GCTCAGCCAGATTCAGTTTCC |
|  | R' | GTCCATGTATGAAGGCTCATGG |
| *CCL5(RANTES)* | F' | CCACAGGTACCATGAAGGTCT |
|  | R' | TTTCGGGTGACAAAGACGAC |
| *TNFRSF10B* | F' | GAGGATTGCGTTGACGAGA |
|  | R' | CCGCGTGCTGATTTATGTGT |
| *CXCL16* | F' | TGACTCAGCCAGGCAAT |
|  | R' | AGCTGGAACCTCGTGTAGTA |
| *IL2RG* | F' | CTGGACTGAACAATCAGTGG |
|  | R' | AGCCAACAGAGATAACCACG |
| *CXCL10* | F' | CTGCCATTCTGATTTGCTGCCT |
|  | R' | CTTGATGGCCTTCGATTCTGGA |
| *IRF3* | F' | GGATTTCGGAATCTTCCAGGCCT |
|  | R' | CTGGGAGTGGGGCCAACACCATG |
| *RIG-I* | F' | TGATCGATTCCATCACTATCCATC |
|  | R' | CCTGTACAATCTCTTCGAATCCTG |
| *MAVS* | F' | GTCACTTCCTGCTGAGA |
|  | R' | TGCTCTGAATTCTCTCCT |
| *IFN-γ* | F' | GAAACGAGATGACTTCGAAAAGCTGACT |
|  | R' | GAGCATCTGACTCCTTTTTCGCTTCCCT |
| *IFN-γR1* | F' | GACGAGCAGGAAGTCGATTATGATCCCGA |
|  | R' | CACCCCGCACATGTAAGACTCCTTCTGCT |
| *IFN-γR2* | F' | CACCGACAGTAAATGGTTCACGGCCGACA |
|  | R' | CACCTCAATGTTTTCTGGCGGCCCGACAGT |
| *HERV-E* | F' | GGTGTCACTACTCAATACAC |
|  | R' | GCAGCCTAGGTCTCTGG |
| *HERV-F* | F' | CCTCCAGTCACAACAACTC |
|  | R' | TATTGAAGAAGGCGGCTGG |
| *HERV-K* | F' | ATTGGCAACACCGTATTCTGCT |
|  | R' | CAGTCAAAATATGGACGGATGGT |
| *ERVL* | F' | ATATCCTGCCTGGATGGGGT |
|  | R' | GAGCTTCTTAGTCCTCCTGTG |
| *ERVMER34* | F' | GAATTCAGTGCCACTAAGCAGAC |
|  | R' | TCGGTATATCCAAGACATGATCC |
| *Syncytin* | F' | TTCACTGCCCACACCCATATG |
|  | R' | CCCCATCAGACATACCAGTTT |
| *STING* | F' | TATCTGCGGCTGATCCTGCCAGCCT |
|  | R' | TGAGACTAGCCTGGGCAACATGG |
| *AIM2* | F' | CCTCTGCTAGTTAAGCTCTCTG |
|  | R' | GCAGTATGTAGTTTGCCTGTGG |
| *HML-2* | F' | AAAGAACCAGCCACCAGG |
|  | R' | CAGTCTGAAAACTTTTCTCTA |
| *HPRT* | F' | CCTCATGGACTAATTATGGAC |
|  | R' | CCCTGTTGACTGGTCATTACA |
