## Supplementary table 3 for "Supraphysiological testosterone induces ferroptosis and activates NF-kappaB mediated immune pathways in prostate cancer through nucleophagy"

Supplementary table 3: List of Antibodies used

| **S No** | **Name** | **Manufacturer** | **Cat No** | **Dilution** |
| --- | --- | --- | --- | --- |
| 1 | Ferritin | Santa Cruz | SC-376594 | 1:1000 |
| 2 | Vinculin (V11F9) | Millipore | Q2925512 | 1:1000 |
| 3 | LC3-Beta | CST | 2775S | WB: 1:1000  IF: 1:100 |
| 4 | NCOA4 (ARA70) | Santa Cruz | SC373739 | 1:1000 |
| 5 | H2AX (JBW301) | Millipore | 05-636 | IF: 1:100 |
| 6 | STING | CST | 13647S | 1:1000 |
| 7 | Actin (AC74) | Sigma | A2228 | 1:1000 |
| 8 | NIK | CST | 4994 | 1:1000 |
| 9 | P100/52 | CST | 4882 | WB: 1:1000  IF: 1:100 |
| 10 | Rel B | CST | 4992T | 1:1000 |
| 11 | TBK1 (D1B4) | CST | 3504S | 1:1000 |
| 12 | TBK1 Phospho | CST | 5483 | 1:1000 |
| 13 | CXCL10 | CST | 14969 | WB: 1:1000  IF: 1:100 |
| 14 | MAVS | CST | 3993 | 1:1000 |
| 15 | p52 mouse monoclonal antibody | Santa Cruz | SC-7386 | 1:1000 |
| 16 | Secondary Mouse for WB | CST | 7076S | 1:5000 |
| 17 | Secondary Rabbit for WB | CST | 7074P2 | 1:5000 |
| 18 | Secondary Mouse- ALEXA FLUOR 488 DYE | Thermo Fisher | A11029 | 1:200 |
| 19 | Secondary Mouse for Confocal | Thermo Fisher | A11034 | 1:200 |
| 20 | Aim2 (D5X7K) | CST | 12948S | 1:1000 |
| 21 | IL-1B (D3A3Z) | CST | 83186S | 1:1000 |
| 22 | IRF7 | CST | 13014S | 1:1000 |
| 23 | p65 | CST | 8242S | WB: 1:1000  IF: 1:100 |
| 24 | MLKL (D216N) | CST | 14993S | 1:1000 |
| 25 | RIP3 (E1Z1D) | CST | 13526S | 1:1000 |
